# A family of archaeal hibernation factors that bind in tandem and protect ribosomes in dormant cells

**DOI:** 10.64898/2026.01.19.700200

**Authors:** Amos J. Nissley, Madison C. Williams, Yekaterina Shulgina, Roan W. Kivimae, Dipti D. Nayak, Jamie H. D. Cate

## Abstract

Under nutrient limitation or stress, ribosome hibernation factors inactivate and protect ribosomes. Although ribosome hibernation plays an important role in microbes, we lack a complete understanding of this process in archaea. Here, we identify a family of hibernation factors, which we designate as single ribosomal subunit inhibitors (SriA-SriD), from the methanogenic archaeon *Methanosarcina acetivorans*. All four *sri* genes are encoded in an operon and each Sri protein inhibits protein synthesis *in vitro*. Deletion of *sri* genes in *M. acetivorans* impaired growth recovery after prolonged stationary phase and also led to depletion of the small ribosomal subunit. Cryo-EM structures show that Sri proteins bind to the ribosome in tandem and form conserved protein-protein interfaces. Sri is broadly distributed across archaeal phyla and *sri* genes frequently co-occur. Together, these findings establish Sri proteins as a distinct group of hibernation factors that protect ribosomes during dormancy and expand our understanding of ribosome hibernation in archaea.

## MAIN

Ribosomes are molecular machines that translate mRNA into protein. Protein synthesis is an energetically demanding process hence under sub-optimal conditions ribosomes are inactivated through ribosome hibernation.^1^ Ribosome hibernation is mediated by proteins, known as hibernation factors, which inhibit translation by occluding functional sites on the ribosome or promoting ribosome dimerization.^2,3^ In addition to suppressing protein synthesis, hibernation factors protect ribosomes from damage by endogenous RNases.^4^ Thus, hibernation preserves an intact pool of ribosomes that remain poised to resume translation once growth conditions become favorable.

Multiple ribosome hibernation factors have been identified in bacteria and eukaryotes. RMF, HPF, and RaiA (previously known as protein Y) are bacterial ribosome hibernation factors that bind to the small ribosomal subunit (SSU) and block mRNA and tRNA binding sites.^5,6^ RMF and HPF also promote the formation of inactive 100S ribosome dimers.^7–9^ An additional bacterial ribosome hibernation factor, Balon, is unique in its ability to bind to actively translating ribosomes.^10^ Similar to their bacterial counterparts, eukaryotic ribosome hibernation factors, such as Stm1/Serbp1^11^, IFRD2^12^, Dapl1^13^, and MDF2^14^ occlude tRNA binding sites on the ribosome. Ribosome dimerization has also been observed in eukaryotes, mediated by the microsporidian hibernation factor MDF1.^15^ Overall, ribosome hibernation factors share the role of preserving ribosomes and halting translation, yet they differ in their structure and modes of ribosome inactivation.

Archaea are major occupants of Earth’s extreme environments including nutrient-poor environments, such as the deep subsurface^16^, where molecular processes that enable dormancy are likely crucial for survival. However, unlike other domains of life, the process of ribosome hibernation is less well understood in archaea. Studies of ribosome inactivation in archaea have mainly focused on factors that dimerize ribosomal subunits. These include archaeal ribosome dimerization factor (aRDF), which dimerizes 30S subunits in the genus *Pyrococcus*^17,18^, and methanogen ribosome dimerization factor (MRDF), which dimerizes 50S subunits in *Methanosarcina acetivorans*.^19^ Previously, we uncovered an archaeal ribosome hibernation factor, **D**ual **r**ibosomal subunit **i**nhibitor (Dri), that contains two ribosome binding lobes, one that targets the large ribosomal subunit (LSU) and the other that targets the small ribosomal subunit (SSU).^20^ While Dri is predominantly found within the Phylum Thermoproteota, we find that genes encoding either one of the two ribosome binding lobes, which are both composed of four cystathionine-β-synthase (CBS) domains, are much more widely distributed in archaea. We designate these gene products as putative **S**ingle **r**ibosomal subunit **i**nhibitors (Sri).

In the methanogenic archaeon *Methanosarcina acetivorans*, Sri proteins are encoded by a four-gene cluster, composed of one N-terminal Dri homolog, one C-terminal Dri homolog, and two additional four tandem CBS domain-containing proteins. We have designated these four proteins **Sri**A-**Sri**D. A mutant *M. acetivorans* strain lacking all four *sri* genes took longer to recover growth after extended stationary phase. Sri proteins protect ribosomes from degradation in extended stationary phase and also inhibit translation *in vitro*. Cryo-EM structures show that Sri proteins bind the *M. acetivorans* ribosome in tandem to occlude functional sites. These results highlight the prevalence of CBS domain-containing proteins as ribosome hibernation factors across archaea, and the potential for their modular use in different physiological contexts and environments.

## RESULTS

### Sri proteins are translation inhibitors that are upregulated under energy limitation

We investigated the role of Sri proteins in the genetically tractable methanogenic archaeon, *M. acetivorans*. In *M. acetivorans*, we identified four putative Sri proteins based on either their homology to Dri or the presence of four tandem CBS-domains (**Fig. 1a-b**). All four Sri proteins are encoded in a gene cluster and we considered all of them as candidate ribosome hibernation factors and labeled them SriA-SriD, according to their gene order. To determine the expression profile of the *sri* genes, we performed global transcriptomics of the wildtype *M. acetivorans* strain (WWM60)^21^ during mid-exponential growth. All *sri* genes are expressed and there is substantial intergenic transcript coverage between each ORF, suggesting that *sriABCD* are under the control of a single promoter (**Fig. 1c**). In contrast, we did not detect transcript reads in the intergenic space between the *sri* operon and the dual CBS domain gene (MA4647) immediately upstream.

**Fig. 1:**
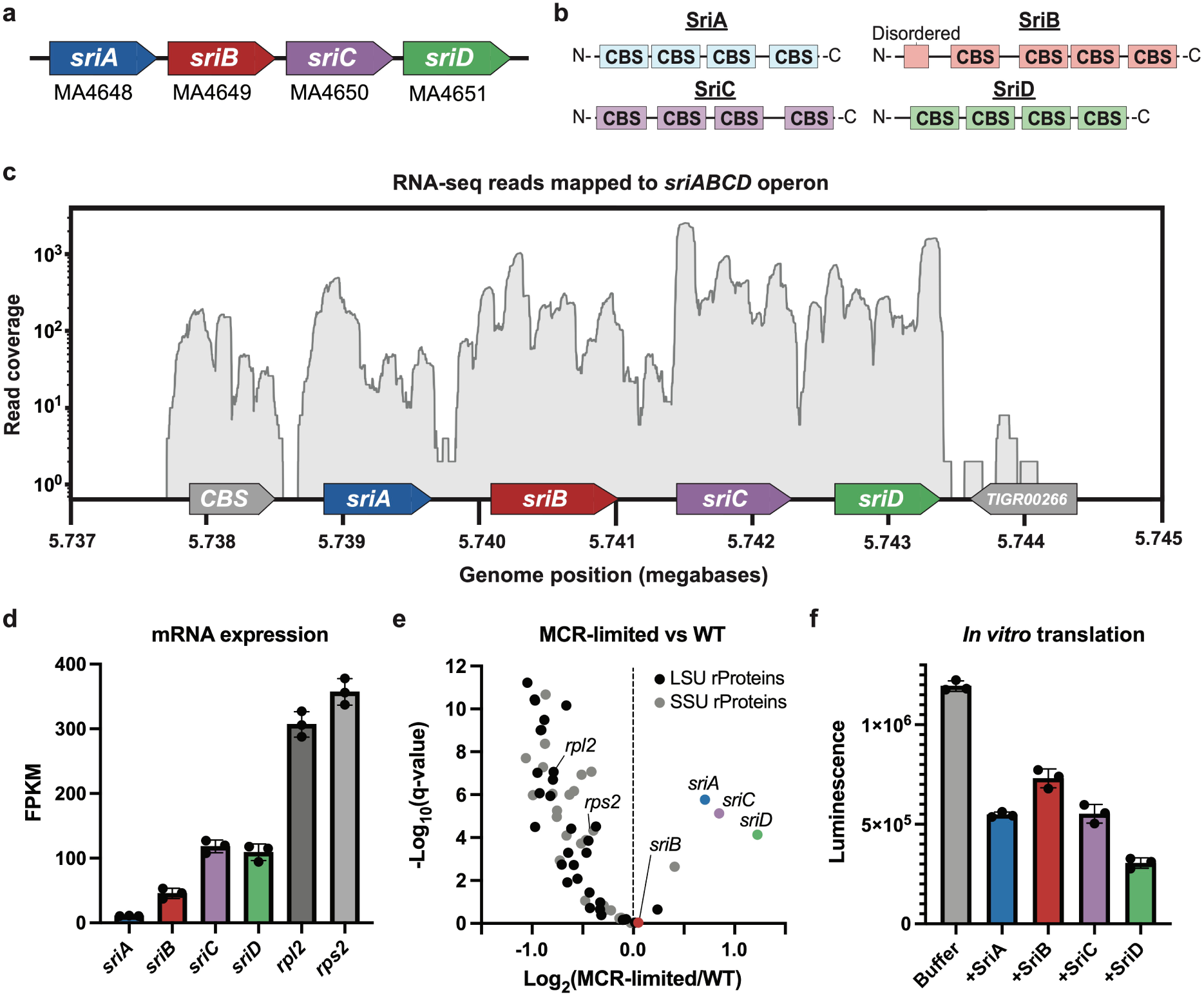
Identification of ribosome hibernation factors in *M. acetivorans*. (a) Gene organization of *sri* genes in *M. acetivorans*. (b) Domain architecture of Sri proteins, which all consist of four tandem cystathionine-β-synthase (CBS) domains. (c) RNA read coverage (forward strand) from exponential phase *M. acetivorans* WWM60 (WT) culture mapped to the *sriABCD* operon. A single representative replicate is shown. (d) mRNA expression levels for Sri and ribosomal proteins from *M. acetivorans* cultures during exponential growth. (e) Differential expression (significance and fold change) in MCR-limited (DDN032 with 0 µg/mL tetracyclin) and WT (WWM60) *M. acetivorans* cultures. (f) *M. acetivorans in vitro* NLuc luciferase translation assay. Sri proteins were added at a final concentration of 6 µM. For the buffer control, 3 µL of Sri buffer A was added to match the buffer of the reactions with added Sri proteins.

Next, we analyzed our previously published transcriptomic datasets to find conditions under which *sriABCD* might be differentially expressed. We found no correlation between *sriABCD* expression and absolute growth rate by comparing *sri* fragments per kilobase of transcript per million mapped reads (FPKM) values from cells cultured in growth medium supplemented with acetate (doubling time, T_D_ ∼80 hr), trimethylamine (T_D_ ∼15 hr), or methanol (T_D_ ∼12 hr).^22,23^ We then asked if the expression of *sri* genes changes if we modulate the energy state of the cell growing on methanol by titrating the cellular concentration of a key metabolic enzyme, methyl-coenzyme M reductase (MCR).^24^ We found that *sri* genes are transcribed during exponential growth in optimal conditions, but have transcript levels lower than ribosomal proteins (rProteins) uL2 and uS2 (**Fig. 1d**). In energy depleted cells, *sriA*, *sriC*, and *sriD* transcripts increase by ∼86% whereas rProtein transcripts are downregulated by ∼30% (**Fig. 1e)**. As MCR induction increases, the upregulation of *sriA*, *sriC*, and *sriD* mRNAs and the downregulation of rProtein mRNAs decreases (**Extended Data Fig. 1a-d**). Anti-correlation between the expression levels of *sri* and rProtein genes specifically under energy limited conditions corroborates their putative role as hibernation factors.

We optimized an *M. acetivorans* lysate-based *in vitro* translation system to test whether Sri proteins inhibit protein synthesis. In the absence of added Sri proteins, we observed robust luminescence from the NLuc reporter (**Fig. 1f**). We recombinantly expressed and purified all four Sri proteins (**Extended Data Fig. 1e**) and the addition of either SriA, SriB, SriC, or SriD to the *in vitro* translation assay decreased luciferase translation. SriD was the most potent inhibitor of the four proteins and its addition to the assay resulted in a 74% reduction in translation (**Fig. 1f**). These results indicate that Sri proteins act as translation inhibitors, consistent with their proposed role in occluding ribosome functional sites during hibernation.

### Sri proteins protect ribosomes in extended stationary phase

In order to query the function of Sri proteins *in vivo*, we generated a markerless quadruple *sri* knockout *M. acetivorans* mutant, Δ*sriABCD*, using our well-established CRISPR editing technology.^25^ We hypothesized that Sri proteins stabilize the ribosome during dormancy, similar to bacterial hibernation factors.^4,26^ To test this hypothesis, we measured the time taken for the Δ*sriABCD* mutant and parental strain, WWM60 (referred to as wildtype or WT), to resume growth after different periods of incubation in stationary phase. Under these circumstances, we expect that the lag time corresponds to the time taken for ribosome synthesis, and would be extended in the absence of hibernation factors like Sri proteins. We grew WT and Δ*sriABCD* starter cultures and subcultured outgrowths at various intervals: mid-exponential, 1 day post-exponential (initial stationary), 19 days post-exponential (stationary), and 28 days post-exponential (extended stationary). We observed no difference in lag time for outgrowths inoculated from mid-exponential or initial stationary phase cultures and only a marginal increase in the lag time for the *ΔsriABCD* stationary phase outgrowths (**Fig. 2a, Supplementary Fig. 1**). However, the Δ*sriABCD* mutant had a 32% longer lag time (74.4 ± 1.8 hr) compared to WT (56.4 ± 2.4 hr) when outgrowths were initiated from extended stationary phase cultures (**Fig. 2b-c**). We did not observe any changes in growth rate or cell yields (measured as maximum optical density) between the two strains under any of the conditions tested (**Supplementary Fig. 1**).

**Fig. 2:**
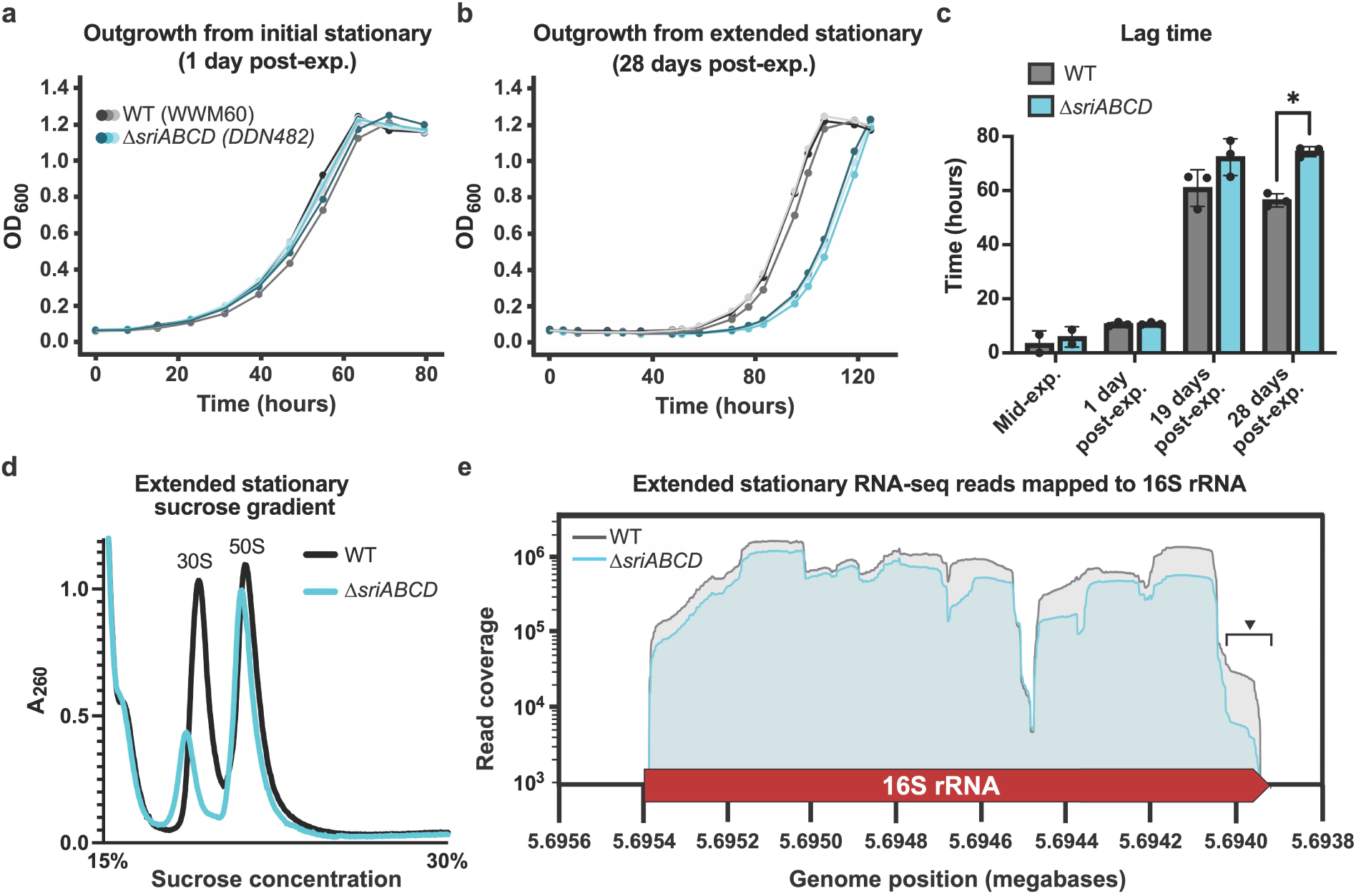
Sri proteins preserve ribosomes *in vivo* during extended periods of stasis. (a-b) Growth curves for outgrowth of *M. acetivorans* WWM60 (WT) and Δ*sriABCD* (DDN482) from (a) initial (1 day post-exponential) and (b) extended (28 days post-exponential) stationary phase. (c) Lag times of outgrowths from mid-exponential phase, 1 day post-exponential, 19 days post-exponential, and 28 days post-exponential. Statistical significance was tested with an unpaired Welch’s t test; * indicates P-value of 8.0x10^-4^. (d) 0.75 mg of protein content from WWM60 (WT) or the Δ*sriABCD* mutant in extended stationary phase was resolved on a dissociating 15-30% sucrose gradient. (e) Paired-end RNA read coverage of *M. acetivorans* 16S rRNA. Carat and bracket indicate the degraded region of the 3′-end that was quantified. One representative technical replicate is shown for WT and Δ*sriABCD*.

To directly test the role of Sri proteins as hibernation factors, we quantified ribosomes in lysates from extended stationary cultures of WT and the Δ*sriABCD* mutant. We passed the lysate through a sucrose gradient under conditions that would dissociate ribosomes into the individual subunits and quantified their abundance. We observed a large decrease in the amount of 30S, but not 50S subunits, from the Δ*sriABCD* mutant compared to WT (**Fig. 2d**). Next, we sequenced total RNA from WT cells in exponential phase as well as WT and Δ*sriABCD* strains in extended stationary phase and aligned reads to the three rRNA loci in the *M. acetivorans* genome. We observe readthrough of the 16S-23S intergenic space in WT at exponential phase, indicative of an immature rRNA pool, but not in the stationary phase samples, which confirms that there is no active transcription of the rRNA under these conditions (**Extended Data Fig. 2a**). In extended stationary phase, we observed a 33% reduction in average 16S rRNA coverage from Δ*sriABCD* samples including an 81% reduction in coverage at the 3′ end, which includes the anti-Shine-Dalgarno (aSD) sequence (**Fig. 2e**). Likewise, when we resolved total RNA on an agarose gel, we could not detect any full-length 16S rRNA in the Δ*sriABCD* extended stationary culture (**Extended Data Fig. 2b**). In contrast to the 16S rRNA, we did not observe a widespread drop in read coverage for the 23S rRNA from the Δ*sriABCD* mutant compared to WT. However, we did observe lower coverage in part of domain IV of the 23S rRNA (helices H61, H64, and H66-H71) from WT at extended stationary phase compared to exponential phase. This region is also more severely degraded in Δ*sriABCD* compared to WT in extended stationary phase (21% reduction) (**Extended Data Fig. 2c**). Taken together, these data suggest that Sri proteins stabilize the ribosome *in vivo* during extended periods of growth stasis.

Additionally, we tested if the role of Sri proteins as hibernation factors extends to stressful conditions. To this end, we exposed WT and the Δ*sriABCD* mutant to a suite of stressors: O_2_ exposure, heat or cold shock, or subinhibitory amounts of the translation inhibitor puromycin. Under all conditions tested, the Δ*sriABCD* mutant grew similarly to WT (**Supplementary Fig. 2**). These data support a context specific role for Sri proteins during periods of growth stasis, i.e. dormancy.

### SriA and SriB bind to the large ribosomal subunit

To understand how Sri proteins mediate ribosome hibernation, we reconstituted Sri-ribosome complexes and determined their structures by cryo-EM. In the structures of the *M. acetivorans* 70S ribosome, we modeled all three rRNAs, 54 annotated rProteins, and a homolog of a previously identified archaea-specific rProtein, aS21^27^ (**Extended Data Fig. 3a**). We also identified density corresponding to a previously undescribed rProtein in the SSU, which we designated aS36. On the SSU, aS36 occupies a position analogous to rProtein eS30, which is found in eukaryotes and some archaea, suggesting that it has a similar structural role (**Extended Data Fig. 3b-c**). The aS36 protein consists of a single domain of unknown function, Pfam DUF5350, and is broadly distributed in members of the phylum Halobacteriota, which includes class II methanogens like *M. acetivorans* (**Extended Data Fig. 3d**). In addition, our high-resolution cryo-EM maps revealed magnesium and zinc binding sites and putative ribosomal RNA (rRNA) methylation sites in the *M. acetivorans* ribosome (**Supplementary Fig. 3**).

In the initial reconstitution, we added all four Sri proteins to *M. acetivorans* lysate and purified ribosomes by RNA affinity purification using poly-lysine (RAPPL).^28^ After focused classification, we determined a 2.0 Å structure of the 70S ribosome (**Supplementary Fig. 4**) with densities in the LSU that correspond to SriA and SriB (**Fig. 3a**). While we resolved the entire SriA protein, only a portion of the SriB N-terminal extension, residues 10-56, were resolved in the cryo-EM reconstruction (**Extended Data Fig. 4a**). The SriB N-terminal extension is predicted to be disordered by AlphaFold^29^, but is ordered while bound to the ribosome in the cryo-EM map. On the LSU, SriA and the SriB N-terminal extension form an interface mediated by the 23S rRNA (**Fig. 3b**). The SriA-SriB interface consists of hydrogen-bonding networks involving ordered water molecules, and the two proteins form a wedge around the conserved peptidyl transferase center (PTC) nucleotide A2617 (2602 in *E. coli*, **Supplementary Table 1**). SriA occupies the A- and P-site tRNA binding sites^30^, and the SriB N-terminal extension interacts with SriA in the A site and extends into the PTC (**Fig. 3c**). Overall, the SriA-SriB complex binds in a position similar to the Dri N-terminal lobe, which is homologous to SriA (**Extended Data Fig. 4b**). Consistent with their role in preserving ribosomes, the 23S rRNA near the SriA and SriB binding sites is more protected from degradation during extended stationary phase in WT compared to the Δ*sriABCD* strain (**Extended Data Fig. 4c-d)**. In the SriA-SriB-ribosome complex, we also observed strong density for the acceptor stem of an E-site tRNA, indicating that SriA binding is compatible with E-site tRNA occupancy (**Supplementary Fig. 5a**). SriA also interacts with a loop in uL16 involved in P-site tRNA binding (**Supplementary Fig. 5b**)^.31,32^

**Fig. 3:**
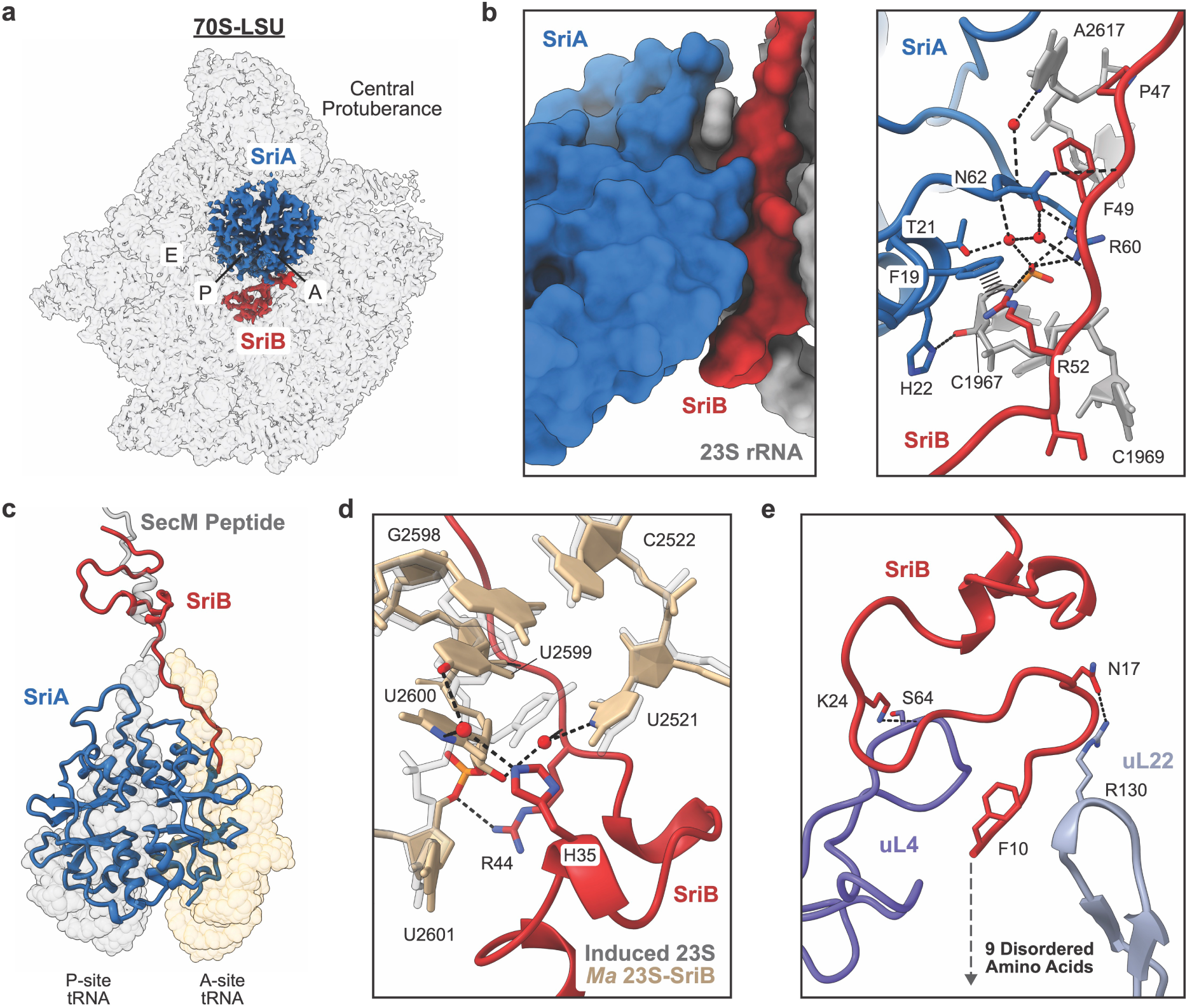
SriA and SriB in complex with the *M. acetivorans* ribosome. (a) Cryo-EM density of SriA and SriB bound to the LSU of the 70S ribosome viewed from the subunit interface. (b) Molecular surface (left) and ribbon (right) representations of the SriA-SriB interface and interactions with the 23S rRNA. (c) Overlay of SriA and SriB with the A-and P-site tRNAs and the SecM peptide from PDB:8QOA^30^. (d) PTC nucleotides involved in the tRNA-induced fit shown with tRNA bound to the *E. coli* ribosome (grey, PDB:8EMM)^34^ or with SriB (red) bound to the *M. acetivorans* LSU (tan). (e) Interaction of SriB with NPET rProteins uL4 and uL22. An additional nine disordered amino acids, not resolved in the structure, extend further down the NPET.

In addition to its interaction with SriA, SriB extends into the PTC and the nascent polypeptide exit tunnel (NPET). In the PTC, SriB induces a conformational rearrangement that is reminiscent of, but distinct from, the active state adopted upon tRNA binding (**Fig. 3d**).^33,34^ The SriB N-terminus also extends past the PTC and interacts with NPET rProteins uL4 and uL22 (**Fig. 3e**). In addition, nine N-terminal amino acids of SriB are unresolved in the cryo-EM map but are oriented further down the NPET. The SriB trajectory down the NPET differs from those of nascent polypeptides and other hibernation factors that target the NPET (**Supplementary Fig. 5c**).

### SriC and SriD bind to the small ribosomal subunit

To identify the binding sites for SriC and SriD, we repeated the Sri-ribosome reconstitutions by adding SriC and SriD to *M. acetivorans* lysate and purifying ribosomes in the absence of exogenous SriA and SriB. After focused classification, we determined a 2.1 Å structure of the 70S ribosome (**Supplementary Fig. 4**) with density in the SSU corresponding to SriD (**Fig. 4a**). SriD occupies the A site of the SSU, where it occludes the mRNA-binding channel (**Fig. 4b**). SriD contains two loops that engage the mRNA decoding center (DC) and interact with conserved 16S rRNA bases involved in the induced-fit mechanism of mRNA decoding (**Fig. 4c**).^35^ The presence of SriD also disrupts inter-subunit bridge B2a, formed between helices H69 of the 23S rRNA and h44 of the 16S rRNA. Upon SriD binding, the apical stem and loop of H69, which normally interacts with h44, becomes disordered (**Extended Data Fig. 5a**). In contrast, SriA binds to the opposite face of H69 in a conformation compatible with bridge B2a formation (**Extended Data Fig. 5b**). Overlay of the SriA and SriD binding sites revealed partial overlap, and the two proteins bind to ribosomes that have distinct SSU rotational states (**Extended Data Fig. 5c-d**). Together, these differences, and the absence of SriA and SriD co-occupancy in our cryo-EM datasets, suggest that SriA and SriD cannot bind to the ribosome simultaneously.

**Fig. 4:**
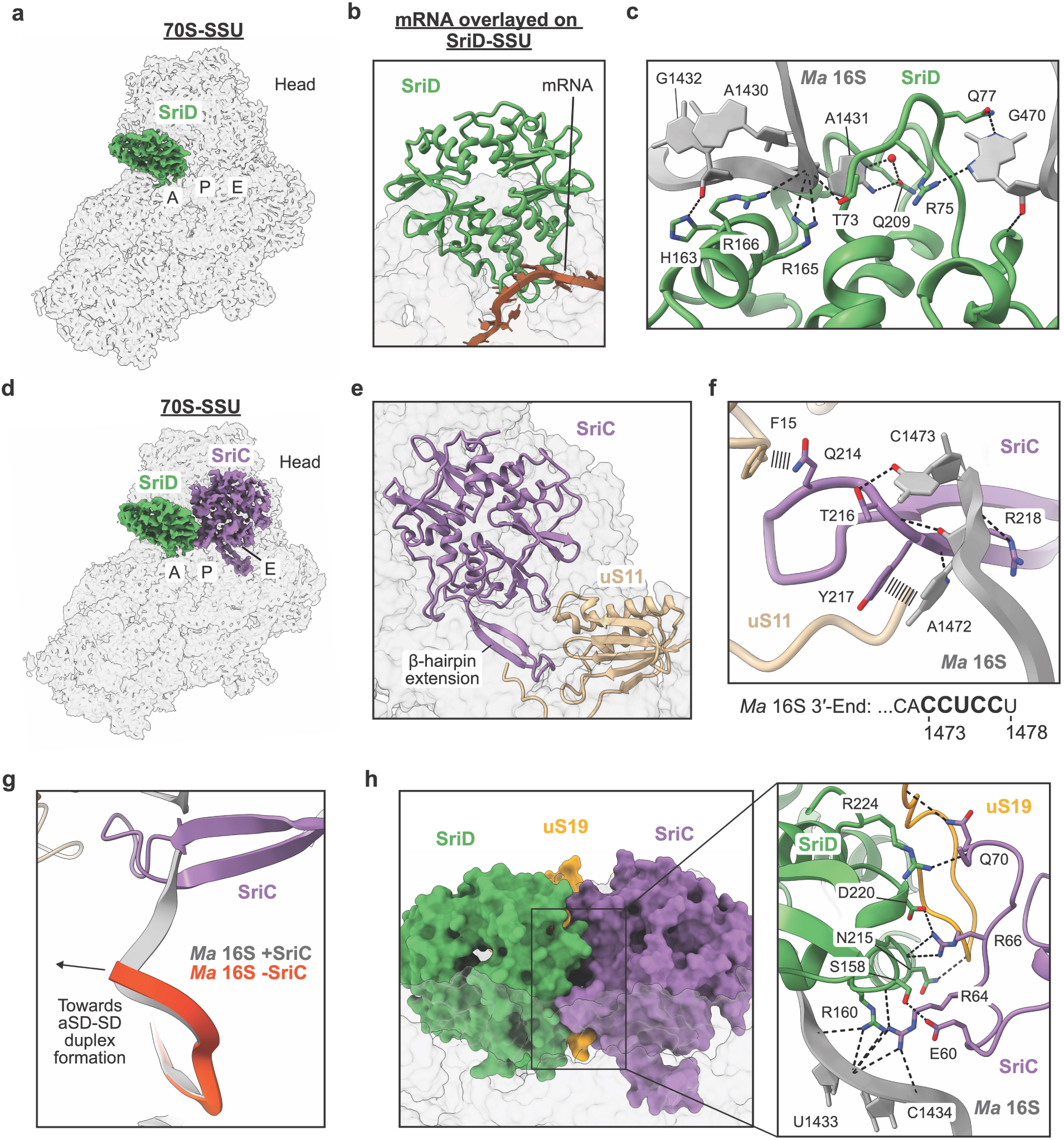
SriC and SriD in complex with the *M. acetivorans* ribosome. (a) Cryo-EM density of SriD bound to the SSU of the 70S ribosome viewed from the subunit interface. (b) Overlay of mRNA bound to the *E. coli* ribosome (PDB:8EMM)^34^ with SriD bound to the *M. acetivorans* ribosome. (c) Interactions between SriD and the mRNA decoding center. (d) Cryo-EM density of SriC and SriD bound to the SSU of the 70S ribosome viewed from the subunit interface. (e) SriC binding in the P- and E-sites of SSU with an extension that interacts with uS11. (f) Interaction between the SriC extension and the 16S rRNA 3′-end. The sequence of the *M. acetivorans* 16S rRNA 3′-end is shown below with the Shine-Dalgarno sequence highlighted in bold. (g) Conformation of the 16S rRNA 3′-end in the presence (grey) or absence (orange) of SriC. The direction of anti-SD (aSD)-SD duplex formation is indicated. (h) Molecular surface (left) and ribbon (right) representations of the SriC-SriD interface and interactions with uS19 and the 16S rRNA.

In our cryo-EM datasets, SriC was detected only in a small portion of particles that also contained SriD. To obtain a structure of SriC bound to the ribosome, we combined particles from the first two datasets and a third dataset (methods) and determined a 2.9 Å structure of the 70S ribosome (**Supplementary Fig. 4**) with density in the SSU corresponding to SriC and SriD (**Fig. 4d**). SriC binds in a similar location to the homologous Dri C-terminal lobe. However, the SriC ribosome-binding loop is distinct from the loop in the Dri C-terminal lobe and forms a β-hairpin extension that engages uS11 (**Fig. 4e and Extended Data Fig. 6a**). The SriC extension also interacts with the 16S rRNA 3′-end, including part of the Shine-Dalgarno (SD) sequence (**Fig. 4f**). This interaction sequesters the 16S rRNA 3′-end in a conformation incompatible with SD-anti SD duplex formation (**Fig. 4g**). In the cryo-EM structure, we observed SriC and SriD binding to the SSU simultaneously. SriC lacks the A-site loop found in the Dri C-terminal lobe, allowing it to form an extensive protein-protein interface with SriD when bound to the SSU (**Fig. 4h**, **Extended Data Fig. 6a**). In addition, SriC and SriD interact with the C-terminal tails of rProteins uS13 and uS19 (**Extended Data Fig. 6b-d**). The uS19 C-terminal tail lies beneath the SriC-SriD interface, where it contacts interface residues (**Fig. 4h**).

### Sri gene clusters are broadly distributed in archaea

To determine the phylogenetic distribution of Sri proteins, we searched 6,968 archaeal species in the Genome Taxonomy Database (GTDB) for *sri* genes based on sequence homology. We assigned homologs of specific *sri* genes by constructing a phylogenetic tree of all Sri homologs and defining subtrees corresponding to each of the four Sri proteins (**Supplementary Fig. 6**). The Dri N-terminal and C-terminal lobe sequences branch with the SriA and SriC clades, respectively, as expected. We found some SriB homologs in multiple archaeal lineages that have lost the N-terminal extension that binds the ribosome (**Supplementary Fig. 7a**, referred to as SriB-short in **Fig. 5a**). We also identified SriC homologs predominantly in the order Nitrososphaerales, many of which have a shorter and less positively charged SSU-binding loop (**Supplementary Fig. 7b**, referred to as SriC-short in **Fig. 5a**). We treat these subsets separately from other *sriB* and *sriC* genes in subsequent analyses, as they may have diverged in function despite their evolutionary relationship. Additionally, we found a widely distributed family of SriA homologs fused with a domain homologous to the bacterial hibernation factors HPF and RaiA that are present in 2,951 (42%) archaeal species. During the preparation of this report, these proteins were shown to function as ribosome hibernation factors and were named Hib.^36^ We also found Hib homologs in some bacteria, predominantly in the phylum Patescibacteriota, class Microgenomatia. In archaea, *hib* and *sri* genes rarely co-occur, with only 91 genomes containing *hib* alongside any *sri* gene (**Fig. 5a-b**). Overall, *sri*, *hib*, or *dri* genes are present in most archaea (4,523 out of 6,968 species, 65%).

**Fig. 5:**
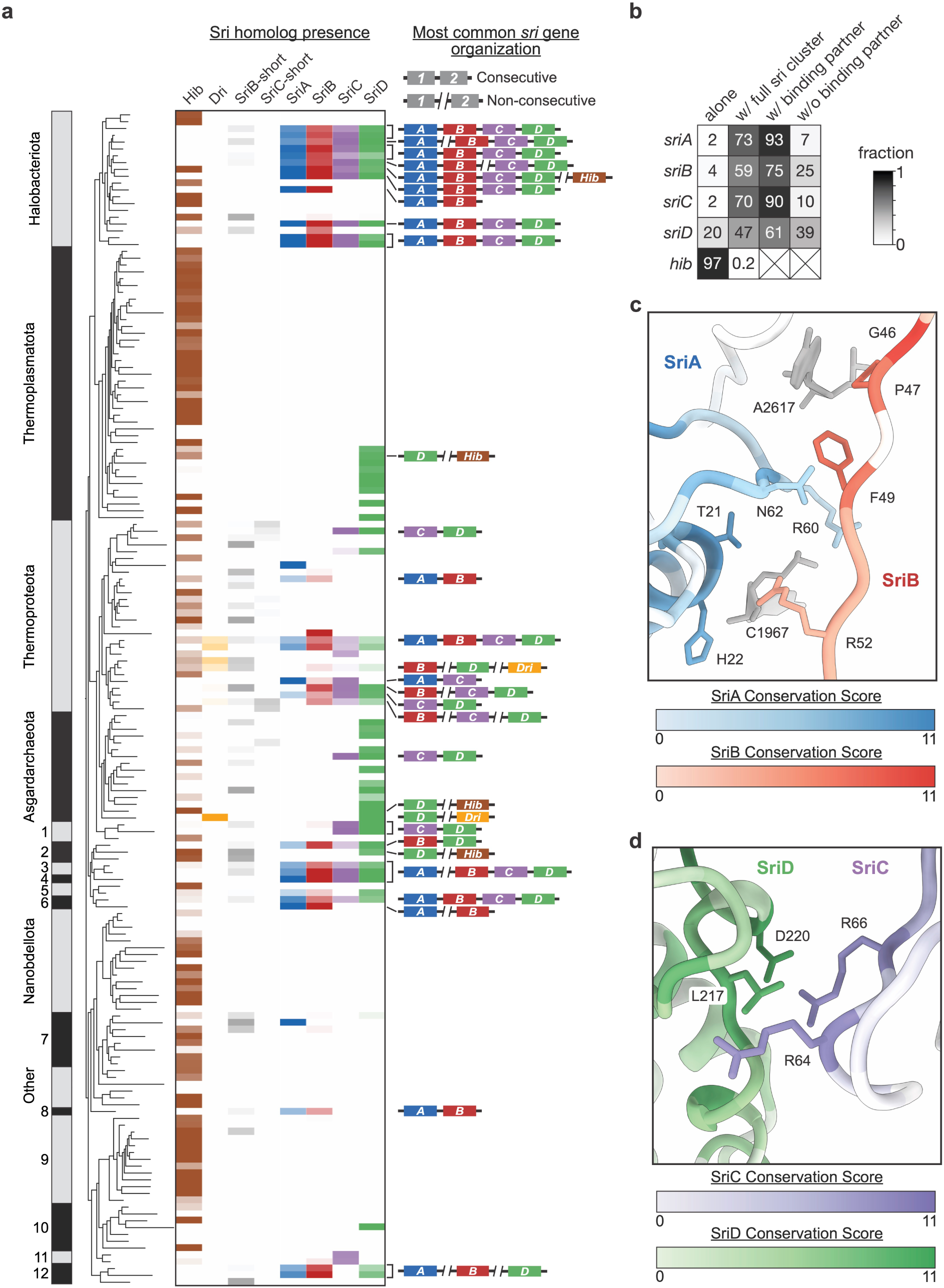
Distribution and conservation of Sri homologs in archaea. (a) Distribution of Sri homologs in GTDB archaea at the order level. Archaeal phyla are indicated on the GTDB tree (left) and Sri homologs are shaded according to occurrence per species in each archaeal order. SriB homologs that lack an N-terminal extension and divergent SriC homologs are shown in the SriB-short and SriC-short columns, respectively. The most common gene organization involving multiple *sri* genes in the indicated orders of archaea is shown on the right. Phylum abbreviations: 1-Korarchaeota, 2-Methanobacteriota, 3-Hydrothermarchaeota, 4-JACRDV01, 5-Methanobacteriota_B, 6-Hadarchaeota, 7-Aenigmatarchaeota, 8-Undinarchaeota, 9-Micrarchaeota, 10-Iainarchaeota, 11-B1Sed10-29, 12-Altiarchaeota. (b) Fraction of Sri and Hib homologs that occur alone (without other *hib* or *sri* genes), co-occur with the full sri cluster (*sriA*, *sriB*, *sriC*, and *sriD*), or co-occur with or without their binding partner (*sriA*-*sriB* and *sriC*-*sriD*). (c) SriA-SriB interface residues are colored based on conservation scores (11 indicates a conserved position and 0 indicates that no physicochemical properties are conserved) from homologs in organisms that encode both proteins. 23S rRNA bases that mediate SriA-SriB interactions are shown in grey. (d) SriC-SriD interface residues are colored based on conservation scores from homologs in organisms that encode both proteins.

The *sri* genes are present in 1,636 (23%) species across 15 archaeal phyla and frequently occur together in a gene cluster, with 1,146 species containing at least two adjacent *sri* genes (**Fig. 5a-b**). The most common gene arrangement is *sriA*-*sriD* in syntenic order (present in 529 species). SriA and SriC frequently co-occur with their binding partners (SriA-SriB and SriC-SriD), while SriB and SriD sometimes occur without their partners (**Fig. 5b**). Several clades of archaea encode a standalone *sriD* gene (**Fig. 5a**), suggesting that SriD may be sufficient to mediate hibernation in these organisms. To evaluate whether Sri homologs bind in pairs to ribosomes across archaea, we examined the conservation of residues^37,38^ involved in Sri-Sri protein contacts on the *M. acetivorans* ribosome. In organisms that encode both SriA and SriB or SriC and SriD, residues at the protein-protein interfaces are conserved (**Fig. 5c-d**). Conversely, in organisms that encode standalone *sri* genes, the residues at the Sri-Sri interfaces are less conserved (**Supplementary Fig. 8**).

## DISCUSSION

Here, we demonstrated that Sri proteins mediate ribosome hibernation in *M. acetivorans*. SriA and SriC bind to the ribosome in similar locations to Dri and the discovery of additional factors SriB and SriD revealed that Sri proteins bind to the ribosome in tandem and inactivate the ribosome. Sri proteins are distinct from other hibernation factors because they form conserved protein-protein interfaces with each other upon binding the ribosome. The coordinated binding of multiple hibernation factors to the ribosome may enhance binding cooperativity or could introduce additional layers of regulation. This mode of binding is reminiscent of some hibernation factors that simultaneously bind to ribosomes with translation elongation factors.^10,13,19,39–42^ SriA-SriB and SriC-SriD complexes cannot bind to the ribosome concurrently because the SriA and SriD binding sites overlap. This suggests that the two sets of proteins target different ribosomes in the cell.

Sri proteins protect the SSU from degradation in extended stationary phase. Depletion of the SSU and degradation of parts of the 16S and 23S rRNA in the Δ*sriABCD* mutant during extended stationary phase leads to an increase in the lag phase upon growth resumption. Therefore, Sri proteins likely help archaea that inhabit nutrient-limited environments recover from dormancy. While ribosome degradation in archaea is not well understood, 3′-5′ RNA degradation is likely mediated by the exosome, though some species in Halobacteriota also encode an RNaseR homolog.^43^ Our cryo-EM structures show that SriC sequesters the 16S rRNA 3′-end and could protect it from exonucleases. *M. acetivorans* 23S rRNA is circularized in the mature LSU^44^, protecting it from exonucleases. This could explain why we observe less depletion of the LSU compared to the SSU in the absence of Sri proteins during extended stationary phase. However, we also observe degradation in domain IV of the 23S rRNA in the absence of Sri proteins, suggesting that there are endonucleases which degrade this region of the LSU.

In *M. acetivorans,* Sri proteins are expressed during active growth, suggesting that they do not bind to actively translating ribosomes, but bind to ribosomes upon translational arrest. Additional proteins or small molecules could regulate Sri binding to ribosomes. CBS-domains often function as metabolite sensors that bind to adenosine nucleotides,^45^ and it remains unknown if nucleotides regulate Sri protein function. Additionally, a purine biosynthesis enzyme, PurH, and translation elongation factor 2, aEF2, dimerize 50S subunits in *M. acetivorans*.^19^ It is unclear how 50S-dimerization or Sri-mediated 70S hibernation are coordinated, or if these pathways can cooperate to mediate ribosome hibernation.

Sri homologs are found across multiple archaeal phyla and represent a broadly distributed set of hibernation factors. While many clades of archaea encode all four Sri proteins, others encode only a subset, suggesting that different combinations or standalone Sri proteins can mediate ribosome hibernation in these organisms. Curiously, the occurrence of *sri* and *hib* (SriA-RaiA/HPF fusion)^36^ genes in archaeal genomes is strongly anti-correlated, implying that these hibernation systems are either redundant or incompatible. Sri, Hib, and Dri hibernation factors have a clear evolutionary relationship through their shared CBS-domain architecture, but the details of the gene fusion and duplication events leading to each protein family remain unclear. Because of their distribution, these factors were likely present in some form in the last archaeal common ancestor. Our work establishes Sri proteins as a widespread set of archaeal hibernation factors and enables further investigation of ribosome hibernation in archaea.

## DATA AVAILABILITY

Ribosome coordinates have been deposited in the Protein Data Bank for the *M. acetivorans* 70S-SriA-SriB complex (9ZNF), 70S-SriD complex (9ZNG), and 70S-SriC-SriD complex (9ZNI). Cryo-EM maps have been deposited in the Electron Microscopy Data Bank for the 70S-SriA-SriB complex (EMD-74446, EMD-74841 EMD-74851, EMD-74853, and EMD-74854), 70S-SriD complex (EMD-74447, EMD-74982, EMD-74984, and EMD-74985), and 70S-SriC-SriD (EMD-74448, EMD-74988, EMD-74989, EMD-74990, and EMD-74992). Raw RNA-seq reads will be deposited in the Sequencing Reads Archive (SRA) under Bioproject number XXXXXXX before publication. Previously published transcriptomic datasets we analyzed can be found in the SRA under Bioproject numbers PRJNA1107246, PRJNA823372, and PRJNA1292878.

## ACKNOWLEDGEMENTS

We thank Dan Toso, Ravindra Thakkar, and Paul Tobias for assistance with cryo-EM data collection (Cal-Cryo). This work was supported by the NSF Center for Genetically Encoded Materials, CHE-2002182. M.C.W. is a National Science Foundation Graduate Research Fellow and Genetic Dissection of Cells and Organisms Training Program trainee (National Institutes of Health T32). Y.S. is a Don Brown Awardee of the Life Sciences Research Foundation. D.D.N. acknowledges funding from the Beckman Young Investigator Award sponsored by the Arnold and Mabel Beckman Foundation, the Alfred P. Sloan Research Fellowship sponsored by the Sloan Foundation, the Simons Foundation Early Career Investigator in Marine Microbial Ecology and Evolution Award, and the Packard Fellowship in Science and Engineering sponsored by the David and Lucille Packard Foundation. D.D.N. is a Chan-Zuckerberg Biohub-San Francisco Investigator.

## Materials and Methods

### *Methanosarcina acetivorans* cultivation, growth measurements, and analysis

*M. acetivorans* was cultured in high salt (HS) media with 50 mM trimethylamine as the sole carbon and energy substrate. The growth media was either buffered with 45 mM NaHCO_3_ and N_2_/CO_2_ (80/20) in the headspace or 50 mM PIPES (pH 6.8) and N_2_ in the headspace^46^. All cultures for growth measurement assays were cultivated in hermetically sealed Balch tubes or anaerobic bottles incubated at 37 °C (Heratherm series incubator, Thermo Fisher Scientific, Waltham, MA, USA). Optical density measurements were taken at 600 nm with a UV-Vis spectrophotometer (Genesys50, Thermo Fisher Scientific, Waltham, MA, USA). For stationary exit experiments, starter cultures were monitored and outgrowths were continuously subcultured at the indicated growth stages/time points from a single starter culture per strain. Starter cultures were incubated at 37 °C during the entire duration of the experiment including the entirety of stationary phase.

Stress challenges were conducted as follows. For O_2_ exposure, cultures were inoculated, grown to OD_600_ ∼0.35-0.4, then treated with 5 mL or 20 mL of ambient air injected via a syringe fitted with a 0.22 µm filter. For temperature shock treatment, cultures were inoculated at 37°C then immediately incubated for 1 hr in ice (tubes placed directly in ice in a covered insulated bucket), 4 °C (in tube rack in unlit room refrigerated to 4 °C) or 45 °C (in tube rack in covered water bath). For puromycin treatment (Cayman Chemical, cat. 13884), sterile, anaerobic stocks were prepared and added to media in Balch tubes at final concentrations indicated prior to inoculation.

Doubling times were calculated by performing a linear regression of log_2_ transformed OD_600_ measurements with the highest R2 values (minimum 5 points). Lag times were calculated by solving the linear regression model for time, x, when y is the log transformed initial OD_600_ value.

### CRISPR-Cas9 plasmid construction and mutant generation

Single guide RNA (sgRNA) sequence (GGCCTGTAAGTATCAAAGAG) for CRISPR-SpyCas9 editing plasmid used to delete *sriABCD* (MA4648-4651) was selected using Geneious Prime v2025.0.3 (https://www.geneious.com) using the CRISPR Finder Tool, as described previously^25^. This guide sequence was introduced via Gibson assembly into *AscI* linearized pDN201 backbone (pMW003). A 1000 bp region of homology upstream and downstream of the MA4648-4651 deletion site was amplified from the *M. acetivorans* genome and introduced into the *PmeI* linearized sgRNA containing vector to form the CRISPR editing plasmid (pMW003) via Gibson assembly. For replication in *M. acetivorans*, the CRISPR editing plasmid was co-integrated with pAMG40 using Gateway BP Clonase II Enzyme mix (Thermo Fisher Scientific, Waltham, MA, USA). Successful co-integration was confirmed by a diagnostic restriction digest with *HindIII.* Sequences of pMW001 and pMW003 were confirmed via Oxford Nanopore sequencing at the Barker DNA sequencing facility at UC Berkeley.

The co-integrated editing plasmid (pMW005) was introduced into a 20 mL culture of *M. acetivorans* WWM60 via liposome mediated transformation as previously described^47^. Transfectants were selected for on plates 1.5% agar medium plates (HS 50mM TMA) with 2 µg/mL puromycin and colonies were screened for the desired chromosomal deletion via PCR. Colonies positive for deletion were grown in liquid media with 2 µg/mL puromycin. Editing plasmid was removed by counterselection on agar plates with 20-80ug/mL 8-aza-2,6-diaminopurine sulfate salt (8-ADP) (Combi-Blocks, Inc., CAS # 65591-11-9). The plasmid cured strain was used for growth experiments in **Supplementary Fig. 2**. All primers, plasmids, and strains used in this study can be found in **Supplementary Tables 2-4**.

### Recombinant expression of Sri proteins

*E. coli* Rosetta (DE3) cells were transformed with a pET-21(+) plasmid containing a Sri protein sequence with a C-terminal His-tag (**Supplementary Table 5**). Cultures of the transformants were diluted 1:100 in 1 L of LB media with 100 μg/mL ampicillin and incubated at 37 °C. Once the OD_600_ reached 0.5, protein expression was induced using 0.5 mM Isopropyl β-D-1-thiogalactopyranosie (IPTG) and the culture was incubated at 37 °C for an additional 2 hours. The cells were then pelleted by centrifugation at 4,000 *xg* for 15 minutes and flash frozen. Cell pellets were thawed and resuspended in Sri lysis buffer (30 mM HEPES-KOH pH 7.5, 500 mM KCl, 20 mM imidazole, and 2 mM DTT) with a Pierce protease inhibitor tablet (ThermoFisher) and lysed by sonication. The lysate was clarified by centrifugation at 25,000 *xg* for 1 hour in an F14-14 x 50cy rotor (ThermoFisher) and then passed through a 0.2 µm filter. A 5 mL His-trap column (Cytiva) was equilibrated with 5 column volumes (CV) of Sri lysis buffer, and the lysate was then loaded onto the column. The column was washed with 10 CV of Sri wash buffer (30 mM HEPES-KOH pH 7.5, 2.5 M KCl, 20 mM imidazole) and the protein was eluted using a gradient from 0-100% Sri elution buffer (30 mM HEPES-KOH pH 7.5, 500 mM KCl, 20 mM imidazole) over 5 CV. Sri proteins were buffer exchanged into Sri buffer A (30 mM HEPES-KOH pH 7.5, 500 mM KCl, 2 mM DTT, 20% glycerol) using PD-10 desalting columns (Cytiva), flash frozen, and stored at -80 °C.

### M. acetivorans in vitro translation

Two 500 mL *M. acetivorans* cultures (1L total) were grown in hermetically sealed anaerobic bottles to mid-exponential phase (OD_600_ of ∼1) and cells were harvested by centrifugation at 4,000 *xg* for 15 minutes. The cell pellet was resuspended in 2.5 mL of *Ma* lysis buffer (50 mM HEPES-KOH pH 7.0, 6 mM MgCl_2_, and 4 mM DTT) per gram of wet cell pellet. 20 µL of RQ1 DNase (Promega) was added and the resuspension was incubated at room temperature for 10 minutes. The sample was then homogenized using a 15G needle and the lysate was clarified using two centrifugation steps at 20,000 *xg* for 20 minutes. The lysate was flash frozen and stored at -80 °C.

NanoLuc luciferase (NLuc) mRNA containing an optimized *M. acetivorans* SD sequence^48^ (**Supplementary Table 6**) was *in vitro* transcribed using T7 RNA polymerase (NEB) and purified using the RNA clean and concentrator kit (Zymo). 30 µL *in vitro* translation reactions were assembled as follows: 15 µL *M. acetivorans* lysate, 2 mM ATP, 2 mM GTP, 2 mM spermidine, 100 µM amino acid mixture (Promega), 900 ng NLuc mRNA, 30 U murine RNase inhibitor (NEB), and an additional 3 mM MgCl_2_ (final concentration of 6 mM Mg^2+^). Either 3 µL of Sri protein or Sri buffer A (control reactions) was added to *in vitro* translation reactions. Reactions were incubated at 37 °C for 30 minutes and then for each reaction, 8 µL was placed in three wells of a 384-well plate. 20 µL of LgBiT buffer (20 mM HEPES pH 7.5, 50 mM KCl, and 10% glycerol) with a 1:50 dilution of NanoGlo substrate (Promega) was added to each well. The plate was incubated for 10 minutes at room temperature and then luminescence was measured in a Spark plate reader (Tecan).

### Sucrose gradients

*M. acetivorans* cultures were grown to stationary phase, maintained for four weeks, and cells were harvested by centrifugation at 4,000 *xg* for 15 minutes. The cell pellet was resuspended in 2.5 mL of *Ma* lysis buffer per gram of wet cell pellet with 20 µL of RQ1 DNase (Promega). The resuspension was incubated at room temperature for 10 minutes, homogenized using a 15G needle, and clarified using two centrifugation steps at 20,000 *xg* for 20 minutes. The protein content of the lysate was measured using a Bradford assay (ThermoFisher). 0.75 mg of lysate protein content was diluted to 150 µL in dissociation buffer (20 mM Tris-HCl pH 7.5, 60 mM NH_4_Cl, 1 mM MgCl_2_, 2 mM DTT) and then loaded on a 15-30% sucrose gradient in dissociation buffer. The sucrose gradients were spun at 41,000 rpm (208,000 *xg*) for three hours in an SW-41 rotor (Beckman-Coulter) and analyzed on a BioComp fractionator.

### RNA sequencing

For all RNA-seq data generated in this study (**Fig. 1c**, **2e**, and **Extended Data Fig. 2**), total RNA was isolated from *M. acetivorans* lysate (as described in the *in vitro* translation and sucrose gradient sections) using the quick RNA miniprep kit (Zymo research) in technical triplicate. The manufacturer protocol was used without the on-column DNase digestion. DNase treatment, cDNA synthesis, library preparation and Illumina Stranded Total RNA sequencing was performed at SeqCenter (Pittsburgh, PA). rRNA depletion step was omitted to enable our downstream rRNA read coverage analyses. Paired-end reads were imported and processed in Geneious Prime v2025.2.2 (insert size 500bp) and aligned to the *M. acetivorans* WWM60 genome using Geneious RNA assembler (configuration settings: Medium-Low Sensitivity; Span annotated mRNA introns; Do not trim). For *sri* operon analysis we resolved read strandedness by aligning paired-end reads of representative sample sets with Bowtie2 v2.3.2. (default parameters) via the Kbase^49^ bioinformatics platform. Strand specific coverage was extracted and calculated from the resulting BAM file using Samtools v1.6. All coverage data can be found in **Supplementary File 1**.

### Ribosome purification and preparation for cryo-EM

To isolate ribosome-Sri complexes for cryo-EM, 180 µL samples were prepared as follows: 90 µL *M. acetivorans* lysate was mixed with 2 mM ATP, 2 mM GTP, 2 mM spermidine, 1 µL of RQ1 DNase (Promega), 180 U murine RNase inhibitor (NEB), and 3 mM MgCl_2_ (final concentration of 6 mM Mg^2+^). For data set one, 6 µM of SriA, SriB, SriC, and SriD were each added to the sample. For data set two, 10 µM SriC and SriD were each added to the sample along with 100 µM amino acid mixture (Promega). For data set three, 10 µM SriC and SriD and 100 µM amino acid mixture (Promega) were added to the sample, but ATP and GTP were not added. The samples were then incubated at 37 °C for 30 minutes. Ribosomes were purified using RAPPL.^28^ After incubation, 220 µL RAPPL Buffer A (25 mM HEPES-KOH pH 7.5, 50 mM KCl, 10 mM MgCl_2_, 2 mM DTT) and 100 µL polylysine magnetic beads (MCLAB) were added to the sample, and the mixture was rotated at room temperature for 20 minutes. The supernatant was then removed from the beads, and the beads were washed three times with 250 µL RAPPL buffer A with a one minute incubation at room temperature in between each wash. To elute ribosomes, the beads were incubated at room temperature for 10 minutes with 50 µL RAPPL elution buffer (RAPPL Buffer A with 2 mg/mL poly-L-glutamic acid (molecular weight 3-15 kDa, Sigma-Aldrich). The elution step was repeated three times. Ribosomes were then concentrated in a 100 kDa molecular weight cut off spin filter (Millipore) and washed with RAPPL Buffer A to remove the poly-L-glutamic acid. *M. acetivorans* ribosomes were quantified using the approximation 1 A_260_=40 µg/mL.

For plunge freezing, *M. acetivorans* ribosomes were diluted to 0.42 mg/mL with RAPPL buffer A. Three-hundred mesh R1.2/1.3 UltrAuFoil grids with a layer of 2 nm amorphous carbon (Quantifoil) were glow discharged in a Pelco easiGlow under a 0.37 mBar vacuum with 25 mA current for 12 seconds. 4 µL of ribosomes was applied to the grid, incubated for three minutes at room temperature, and plunge-frozen in liquid ethane using a Vitrobot Mark IV at 4 °C with 100% humidity.

### Cryo-EM data acquisition and image processing

Cryo-EM movies were collected with a 300 kV Titan Krios microscope with a BIO-energy filter and Gatan K3 camera (**Supplementary Table 7**). Data was collected with a super-resolution pixel size of 0.41 Å (physical pixel size of 0.8293) over a defocus range of -0.5 to -1.5 µm with a total electron dose of 40 e^-^/Å^2^ split over 40 frames. Data collection was automated with Serial-EM.^50^

Cryo-EM image processing was done in cryoSPARC 4^51^ (**Supplementary Fig. 9**). Movies were binned to the physical pixel size and patch motion corrected. The CTF of micrographs were estimated using patch CTF estimation and micrographs with poor CTF fit were manually excluded from further processing. Particles were picked using template picking with ribosome 2D templates generated in cryoSPARC. Particles were extracted with a box size of 512 pixels and cropped to 1/8 of the full box size. 2D classification was performed with 100 classes and 2D classes containing features consistent with ice or the foil edge were excluded. The particles were then subjected to heterogeneous refinement with 12 classes and classes consistent with 70S ribosomes were selected for further processing. 70S particles were re-extracted at the full box size and were aligned using homogeneous refinement with the per-particle defocus optimization, per-group CTF parameter optimization, and minimization over per-particle scale options selected. Local motion was then corrected using reference-based motion correction in cryoSPARC and the particles were refined a second time.

For dataset one, 70S particles containing SriA and SriB were classified using 3D classification with a mask on SriA and 3 classes. The class with SriA and SriB density was then refined. For data set two, 70S particles were refined locally on the SSU, and particles containing SriD were classified using 3D classification with a mask on SriD and 3 classes. The class with SriD density was then subjected to a second round of 3D classification using a mask on SriD and 2 classes. The class with density for all SriD domains was then refined. For data set three, 70S particles were classified based on SriD occupancy as described above. SriD containing particles from data sets one and two were then combined and particles also containing SriC were classified using 3D classification with a mask on SriC and 10 classes. SriD containing particles from data set three were also classified on SriC occupancy. Particles from all three datasets containing density for SriC and SriD were then combined and further classified using 3D classification with a mask on SriC and 3 classes. The class with the best density for SriC was then further refined.

After refinement, we used the ‘Fit in Map’ tool in ChimeraX^52^ to calibrate the magnification of the cryo-EM map using the X-ray crystal structure of the *Haloarcula marismortui* 50S subunit^53^, resulting in a pixel size of 0.8264 Å. Local resolution was calculated in Relion 4^54^ using the Relion implementation. For figure presentation, cryo-EM maps were supersampled using the ‘make a smoother copy’ function in COOT.^55^

### Model building

Composite maps were generated for the 70S-Sri complexes. For the 70S-SriA-SriB and 70S-SriC-SriD complexes, we aligned the SSU-focused, SSU head-focused, and LSU-focused refined maps to the global 70S ribosome map using the ‘Fit in Map’ tool in ChimeraX.^52^ For the 70S-SriD complex, we aligned the SSU-focused, and LSU-focused refined maps to the global 70S ribosome map. The focus refined maps were then resampled using ‘vop resample’, scaled according to their standard deviations from the ‘Map statistics’ tool, and added using ‘vop add’.

Initial models were built using ModelAngelo^56^ provided with sequences for *M. acetivorans* rRNAs and rProteins. The linear *M. acetivorans* 23S rRNA sequence was used for modeling to maintain consistency with nucleotide numbering and because the circularized H1 extension in *M. acetivorans* 23S rRNA^44^ is disordered in our cryo-EM maps. Unknown proteins were built using ModelAngelo without provided sequences. Protein models were inspected for completeness and manually built in regions where the model was incomplete. RNA models were manually inspected at each nucleotide for correct modeling and post-transcriptional modifications. For the SriA-SriB and SriD bound ribosomes, there was strong density for the acceptor stem of an E-site tRNA, which was modeled using the sequence from *M. acetivorans* tRNA^Phe^. Magnesium ions were identified using the ‘unmodeled blobs’ tool in COOT^55^ and modeled manually. No attempt was made to systematically model potassium ions. The LSU and SSU models were then refined in their individual focused refined maps using Phenix real-space refinement.^57^ Manual adjustments were made to the models in COOT^55^ and ISOLDE^58^ and the resulting models were docked into the composite map. For the SriA-SriB and SriD bound ribosomes, phenix.douse^57^ was used to model solvent using the default settings in the individual sharpened focused refined maps and the solvent models were docked into the composite maps. The final models were validated in Phenix (**Supplementary Table 8**).

### Bioinformatic analysis

To determine the phylogenetic distribution of aS36, we used the homology search tool HMMER 3.4^59^ to search the predicted proteomes of archaeal representative species in the Genome Taxonomy Database (GTDB, version 226.0)^60^. First, we built a profile HMM built from the *M. acetivorans* aS36 sequence using the HMMER hmmbuild program and then used the HMMER program hmmsearch to query all proteomes, taking all resulting hits with e-value < 1e-5 as homologs of aS36. We aligned these sequences using mafft v7.525^61^ with default settings and built another profile HMM to search again but no additional hits with e-value < 1e-5 were found. The final list of sequences can be found in **Supplementary Files 2-3**.

To search for Dri homologs, we aligned the 92 Dri hits found in Nissley et al (2025)^20^ using mafft with default settings and built a profile HMM using the HMMER hmmbuild program. After searching archaeal GTDB representative proteomes using the HMMER program hmmsearch, hits with e-value < 1e-100 and protein length greater than 500aa were used as Dri sequences for visualization on the Sri protein tree. These Dri homologs were aligned using mafft and split into N- and C-lobes, respectively, based on positions 14-303 and 324-640 in the *Pyrobaculum calidifontis* sequence.

To search for SriA, SriB, SriC, and SriD homologs, we built initial profile HMMs from the *M. acetivorans* sequences. After an initial search of the archaeal GTDB representative proteomes, we identified a subset of genomes where the best hits for the four models were arranged sequentially as four adjacent genes (SriA-SriB-SriC-SriD). These SriA, SriB, SriC, and SriD sequences were aligned separately using mafft and used to build another set of profile HMMs. After an additional round of searches with the final models, all hits with e-value < 1e-35 and protein lengths between 100 and 500 amino acids were pooled. These protein sequences were used to construct a Sri homolog tree, also including the Dri N- and C-lobe sequences. Sequences were aligned using mafft and columns with more than 99% gaps were removed from the alignment. A phylogenetic tree was built using IQTREE v3.0.1^62^ using LG model and default settings. Phylogenetic trees were visualized using the Interactive Tree of Life (iTOL v6)^63^ webserver.

Upon building the tree, we identified a protein clade corresponding to a fusion of SriA and a RaiA/HPF homolog, now known as Hib^36^. Since Hib homologs hit the SriA model weakly, we separately searched for Hib homologs by building a profile HMM from sequences corresponding to this subclade on the tree, searching all GTDB representative proteomes, and including all hits with e-value <1e-35 into the final version of the tree (**Supplementary Fig. 6**).

We defined subtrees corresponding to each of the Sri genes where sequences predominantly hit the appropriate Sri model with the most significant e-value. All sequences falling within each subtree were considered homologs of the respective Sri gene. These subtrees are colored in **Supplementary Fig. 6**, information on the protein sequences can be found in **Supplementary File 4**, and sequences for the assigned Sri and Hib genes can be found in **Supplementary Files 5-9**. We further examined some Sri homologs for sequence features unique to their mode of ribosome binding. For SriB, the length of the N-terminal extension was assessed by aligning each homolog and the *M. acetivorans* SriB sequence to the SriB profile HMM using the HMMER program hmmalign and assessing the length of the homolog’s sequence that aligns to positions 27-56 in *M. acetivorans* SriB, the conserved region of the extension. SriB homologs with N-terminal extension length <= 15 were categorized as SriB-short (**Supplementary Fig. 7a**). For SriC, we assessed the length and electrostatic charge of the ribosome binding loop by aligning each homolog and the *M. acetivorans* SriC sequence to the SriC profile HMM and assessing the length and charge of the homolog’s sequence that aligns to positions 191-225 in *M. acetivorans* SriC. A clade in the SriC subtree, primarily representing a diversification in the order Nitrososphaerales with less positive ribosome binding loop, was dubbed SriC-short and considered separately (**Supplementary Fig. 7b**). In subsequent analyses of conservation, co-occurrence, and phylogenetic distribution, SriB-short and SriC-short sequences were omitted.

We looked for Sri and Hib homologs in the GTDB bacterial species using a similar approach. Briefly, we searched the predicted proteomes of the GTDB representative bacterial species using the Sri and Hib profile HMMs as described above, keeping hits spanning over 75% of the model length with e-value < 1e-15 and protein length < 500, and building a protein tree together with the archaeal Sri, Dri, and Hib sequences as described above. Most bacterial hits clustered outside the Sri and Hib subclades, likely representing other CBS-domain-containing proteins, except for a single clade of 146 bacterial sequences that branched within the Hib subtree. More information on these sequences can be found in **Supplementary Files 4 and 10**.

To subset archaeal Sri hits based on co-occurrence with Sri genes for conservation analysis of the protein interfaces, we first selected the respective Sri gene with the most significant e-value if there were multiple hits. For SriB, we only considered sequences with long N-terminal extensions (>15aa aligning to the region specified above). Sequences were aligned using mafft and conservation scores^37^ were calculated in Jalview^38^.

---

**Extended Data Fig. 1:**
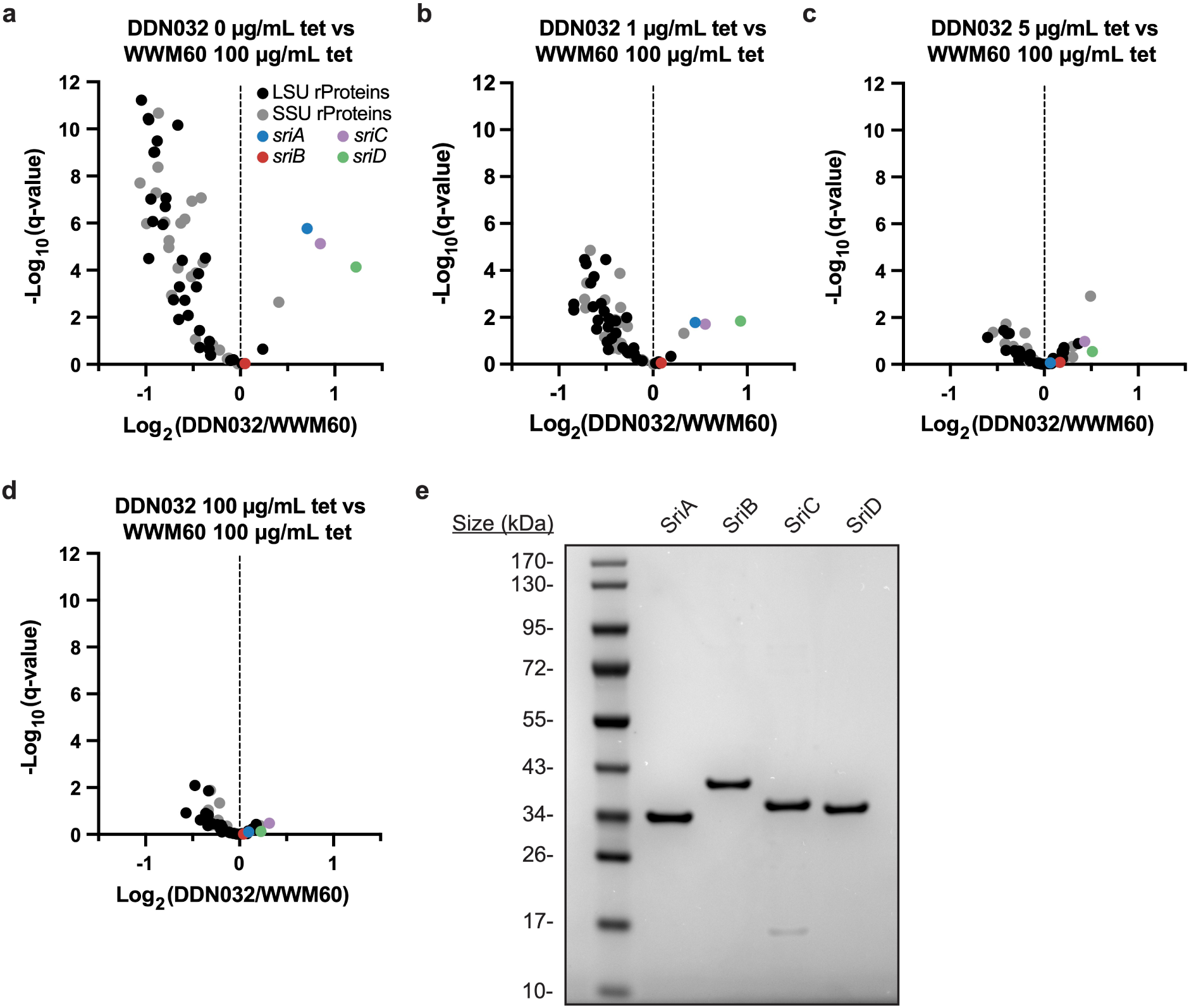
Differential expression of *sri* transcripts under Methyl-Coenzyme M Reductase (MCR) limitation and Sri protein purification. (a-d) Differential mRNA expression (significance and fold change) in DDN032 (tetracycline inducible MCR expression) and WWM60 (WT) *M. acetivorans* cultures. MCR expression was varied across panels a-d with increasing concentrations of tetracycline (tet). Panel a is reproduced from fig. 1 for comparison. (e) Recombinantly expressed and purified Sri proteins resolved on a 4-12% polyacrylamide gel stained with Coomassie brilliant blue.

**Extended Data Fig. 2:**
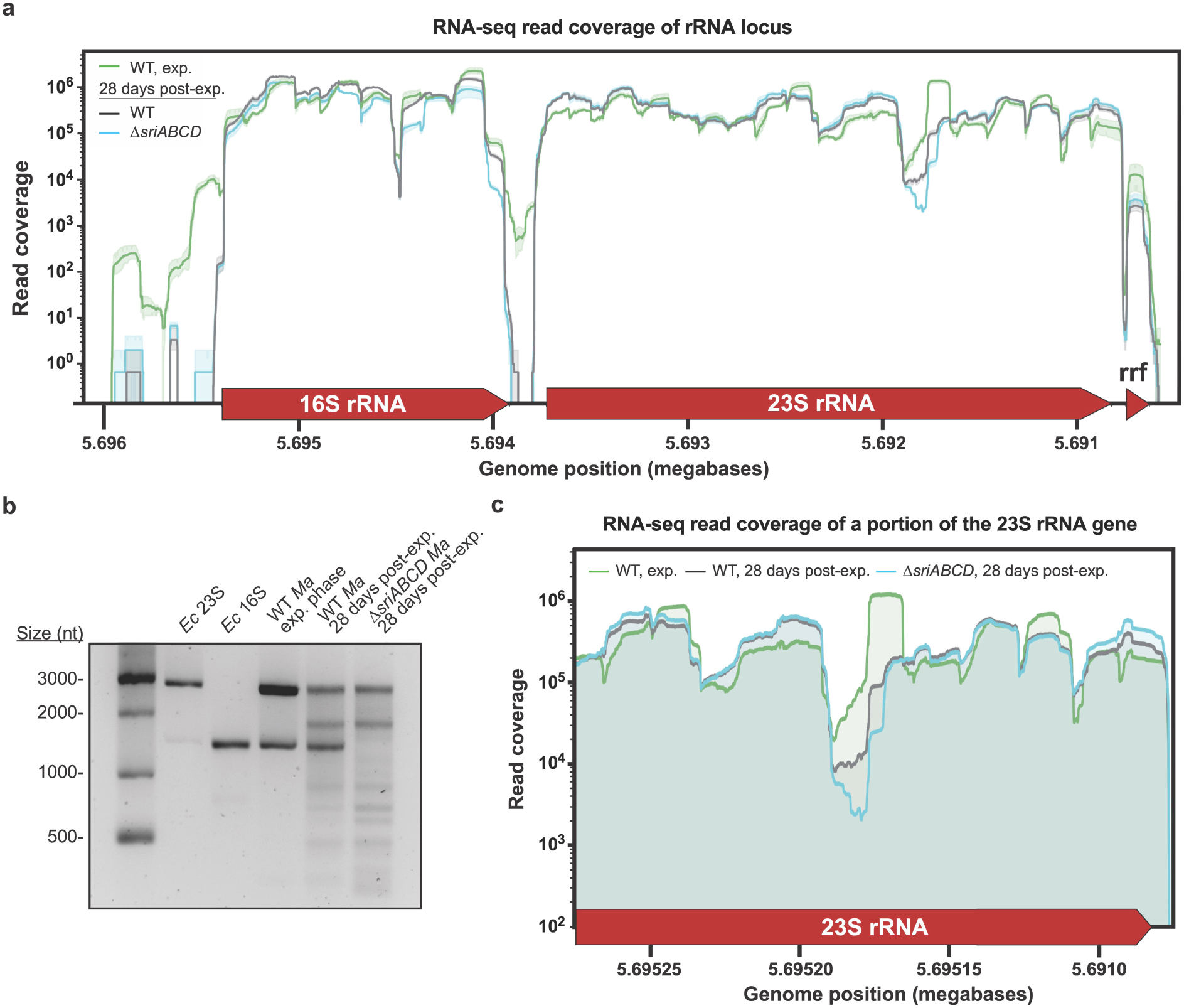
Analysis of *M. acetivorans* rRNAs from exponential and stationary phase. (a) Total RNA read coverage of an rRNA locus. The mean of 3 technical replicates per condition (strains and growth phases) is shown as a solid line and shading indicates the 95% confidence interval. (b) rRNAs resolved on a 2% agarose gel. Purified *E. coli* (*Ec*) 23S and 16S rRNAs serve as standards in the left two lanes. Total RNA from *M. acetivorans* (*Ma*) cultures at exponential phase or 28 days post exponential (post-exp.) phase were isolated and resolved in the right three lanes. (c) RNA-seq read coverage for a portion of the 23S rRNA gene. One representative replicate per strain is shown for visual clarity.

**Extended Data Fig. 3:**
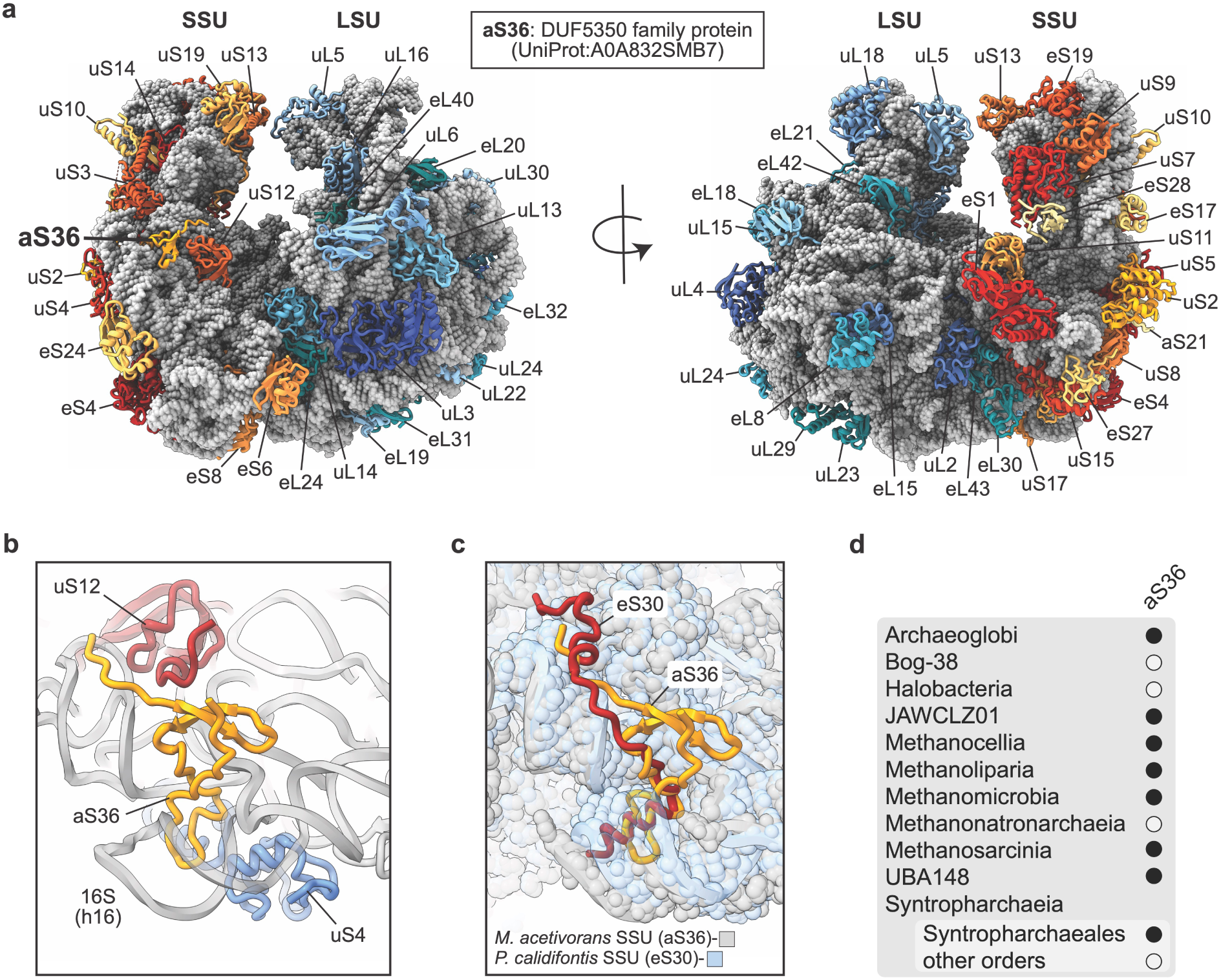
rProteins in the *M. acetivorans* ribosome. (a) rProteins are labeled on the model of the *M. acetivorans* 70S ribosome. The aS36 label is bolded and its previous annotation, including the accession code, is shown in the middle box. (b) aS36 is in proximity to uS4 and uS12 and interacts with 16S rRNA helix h16. (c) Overlay of the SSUs from *M. acetivorans* (grey) and *P. calidifontis* (blue, PDB:9E71^20^) showing the locations of aS36 and eS30, respectively. (d) Distribution of aS36 across the archaeal phylum Halobacteriota. Closed and open circles represent clades where aS36 homologs were or were not identified, respectively.

**Extended Data Fig. 4:**
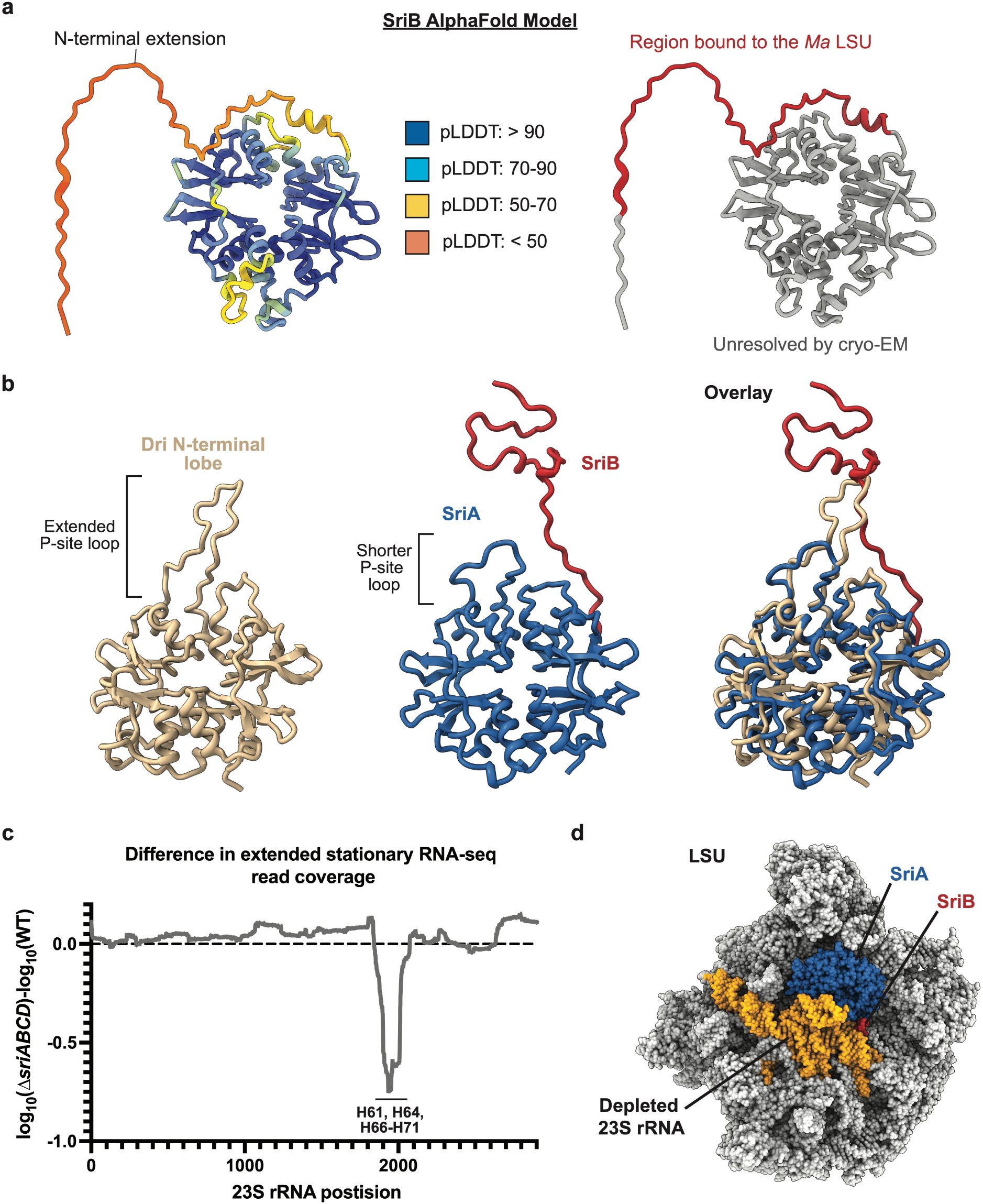
Structural features of SriA and SriB. (a) AlphaFold^29^ prediction of SriB colored by (left) confidence score or (right) regions that are ordered (red) or disordered (grey) in the cryo-EM structure of SriB bound to the *M. acetivorans* ribosome. (b) Comparison of the Dri N-terminal lobe (left) with the SriA-SriB complex (middle). (right) Alignment of the *M. acetivorans* LSU in complex with SriA and SriB to the *P. calidifontis* LSU in complex with Dri (PDB:9E6Q^20^), highlighting the overlap in Sri and Dri binding sites. (c) Difference in the log_10_ RNA-seq read coverage for the 23S rRNA from Δ*sriABCD* (DDN482) and WWM60 (WT) at extended stationary phase. 23S rRNA helices with depleted read coverage in the Δ*sriABCD* mutant are labeled. (d) 23S rRNA positions with depleted RNA-seq read coverage in Δ*sriABCD* at extended stationary phase are shown in yellow and the SriA and SriB binding sites are shown in blue and red, respectively.

**Extended Data Fig. 5:**
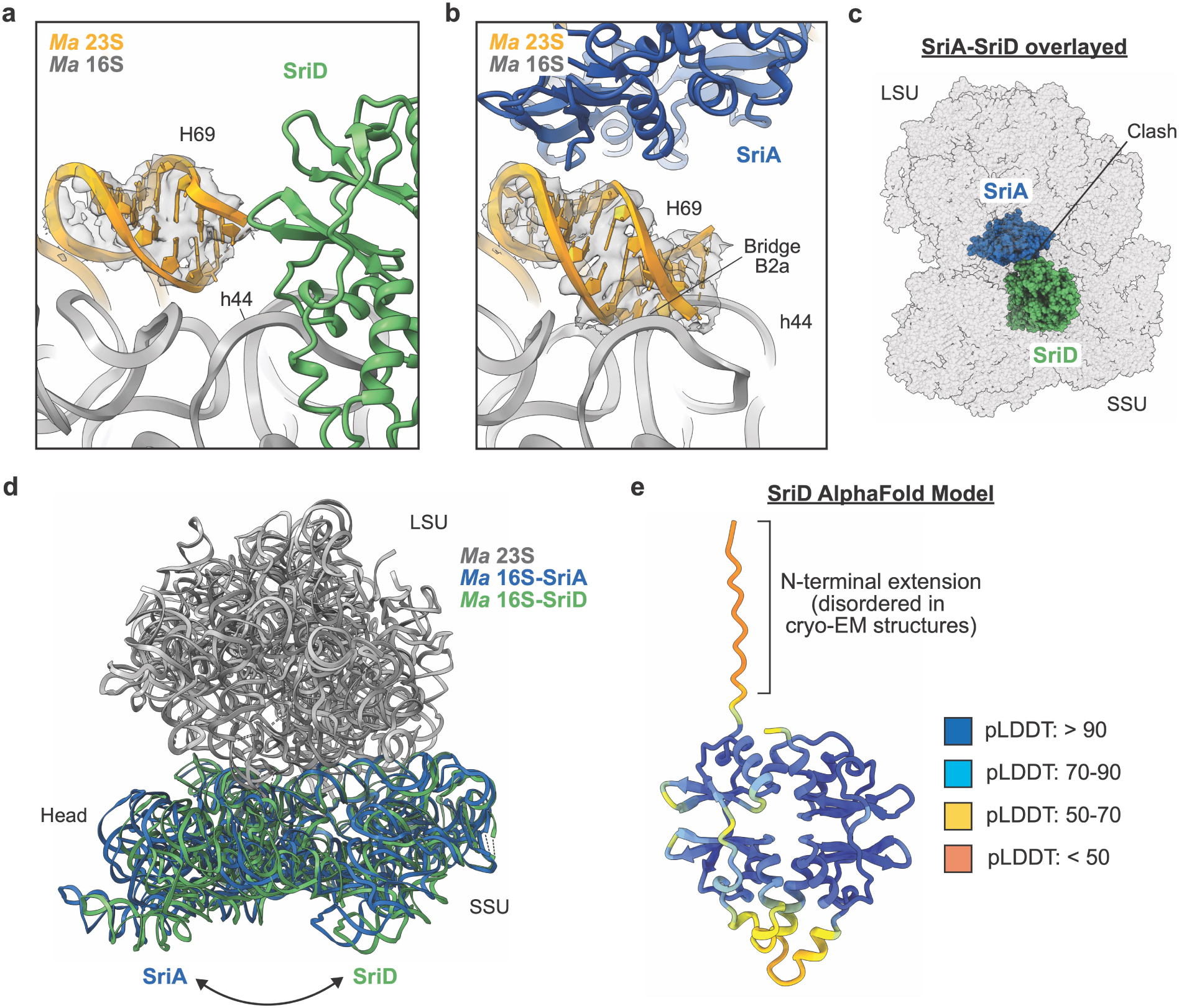
Differences in SriA and SriD binding to the ribosome. (a-b) Conformations of LSU helix H69 and SSU helix h44 in the presence of (a) SriD or (b) SriA. Bridge B2a is formed during SriA binding but is absent in the presence of SriD. (c) Overlay of SriA and SriD binding sites on the ribosome. Simultaneous binding of SriA and SriD would result in a steric clash between the two proteins. (d) *M. acetivorans* ribosomes bound to SriA (blue) and SriD (green) aligned to their 23S rRNA (grey) to highlight differences in the rotational state of the SSU. (e) AlphaFold prediction of SriD colored by confidence score. The SriD N-terminal extension is not resolved in the cryo-EM structure of SriD bound to the *M. acetivorans* ribosome.

**Extended Data Fig. 6:**
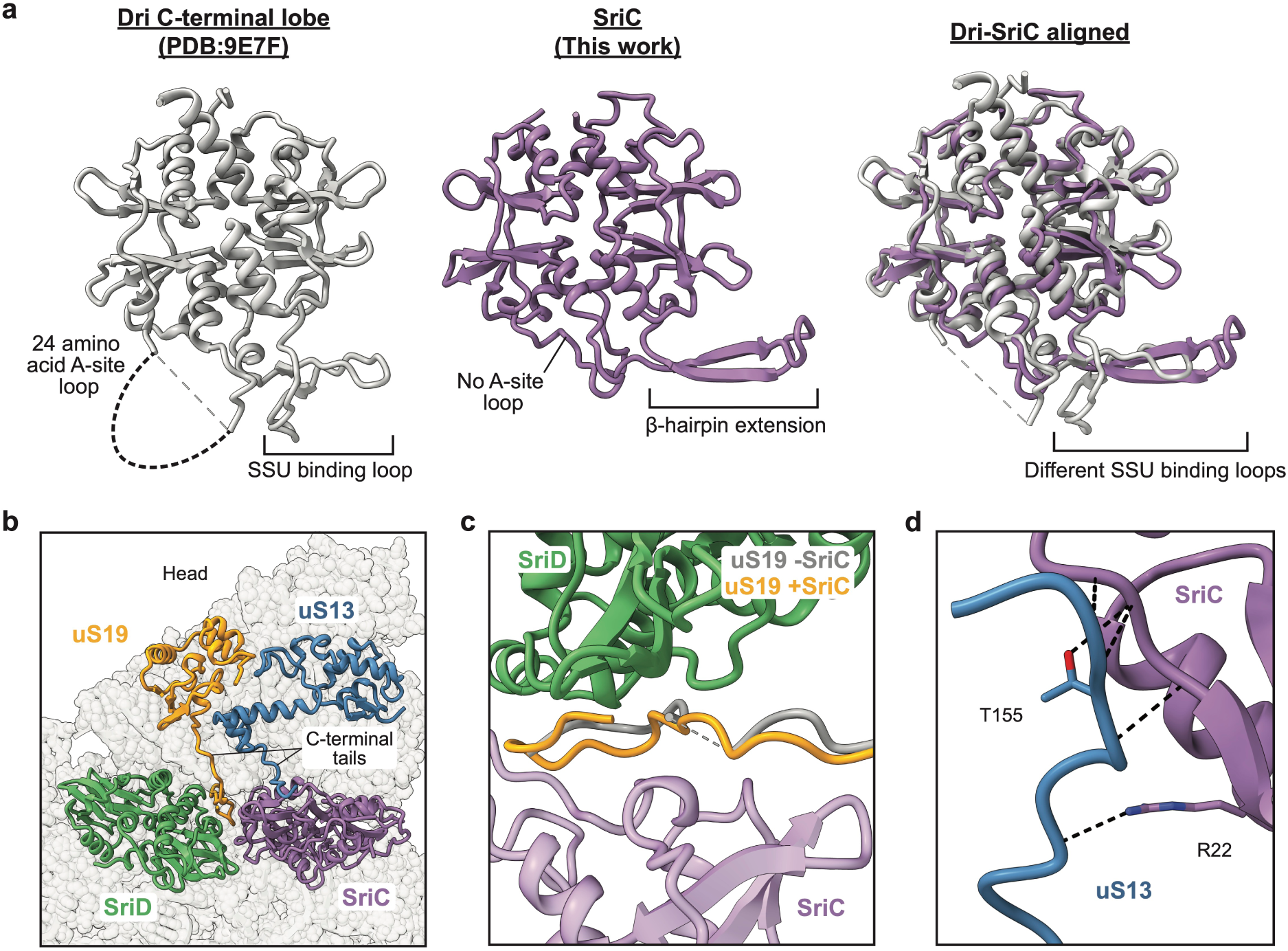
SriC and SriD interactions with the SSU. (a) Comparison of (left) the Dri C-terminal lobe (PDB:9E7F^20^) and (middle) SriC. (right) Alignment of the Dri C-terminal lobe with SriC. (b) Interactions of SriC and SriD with the C-terminal tails of uS13 and uS19. (c) Conformation of the uS19 C-terminal tail in the presence of SriD (grey) or of both SriC and SriD (yellow). (d) Interactions between SriC and the uS13 C-terminal tail.

