## Supplemental Figures and Tables for "A family of archaeal hibernation factors that bind in tandem and protect ribosomes in dormant cells"

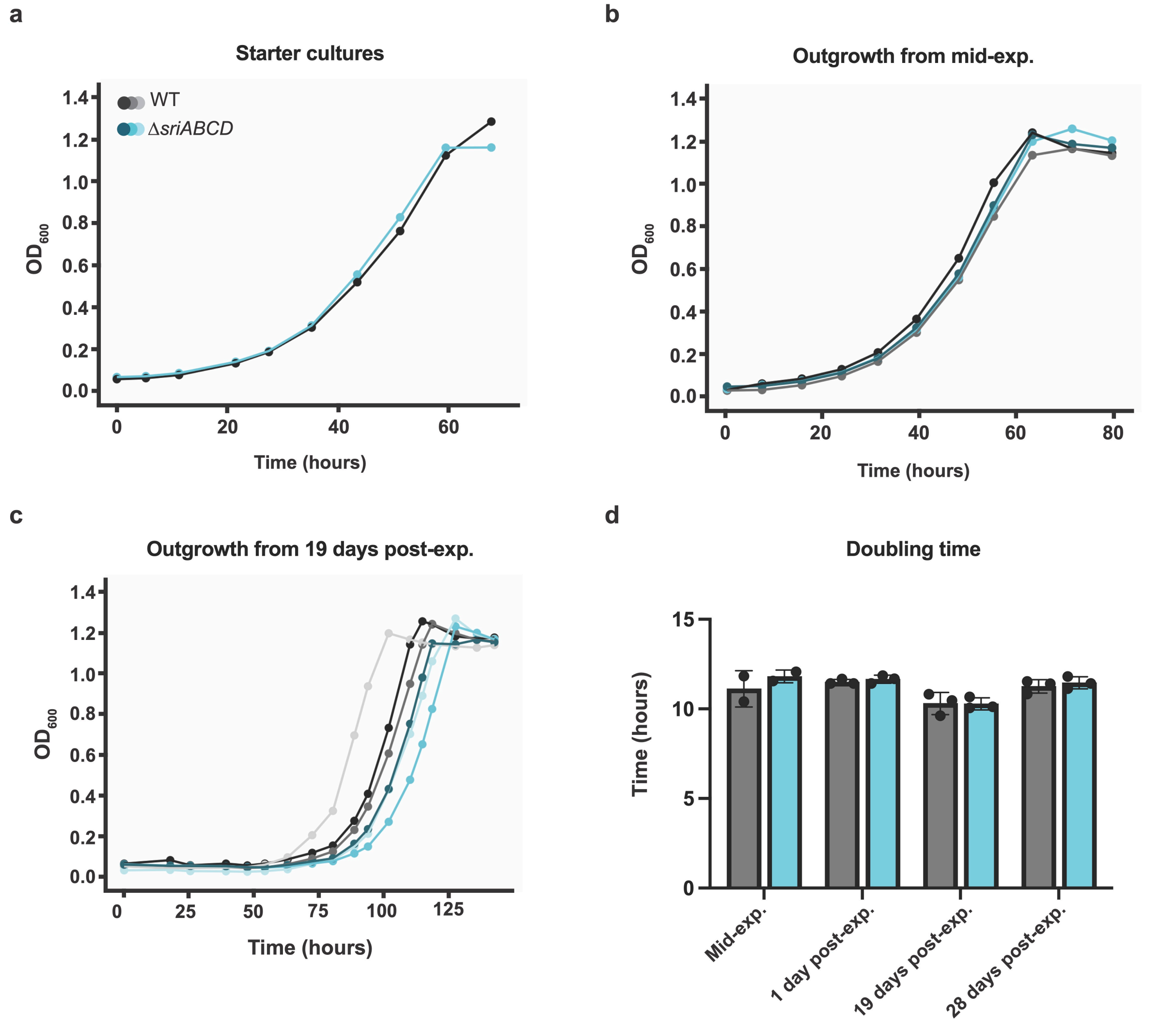


**Supplementary Fig. 1:** Additional stationary phase recovery growth curves. (a-c) Starter cultures, mid-exponential (mid-exp.) and 19 days post-exponential (post-exp.) outgrowth curves. (d) Doubling times from all outgrowths; error bars represent standard deviation. Gray: WT - WWM60, parent strain; blue: Δ*sriABCD* - DDN482, harbors editing plasmid (pMW005). Mid-exp outgrowths cultured in duplicate, all other outgrowths cultured in triplicate.

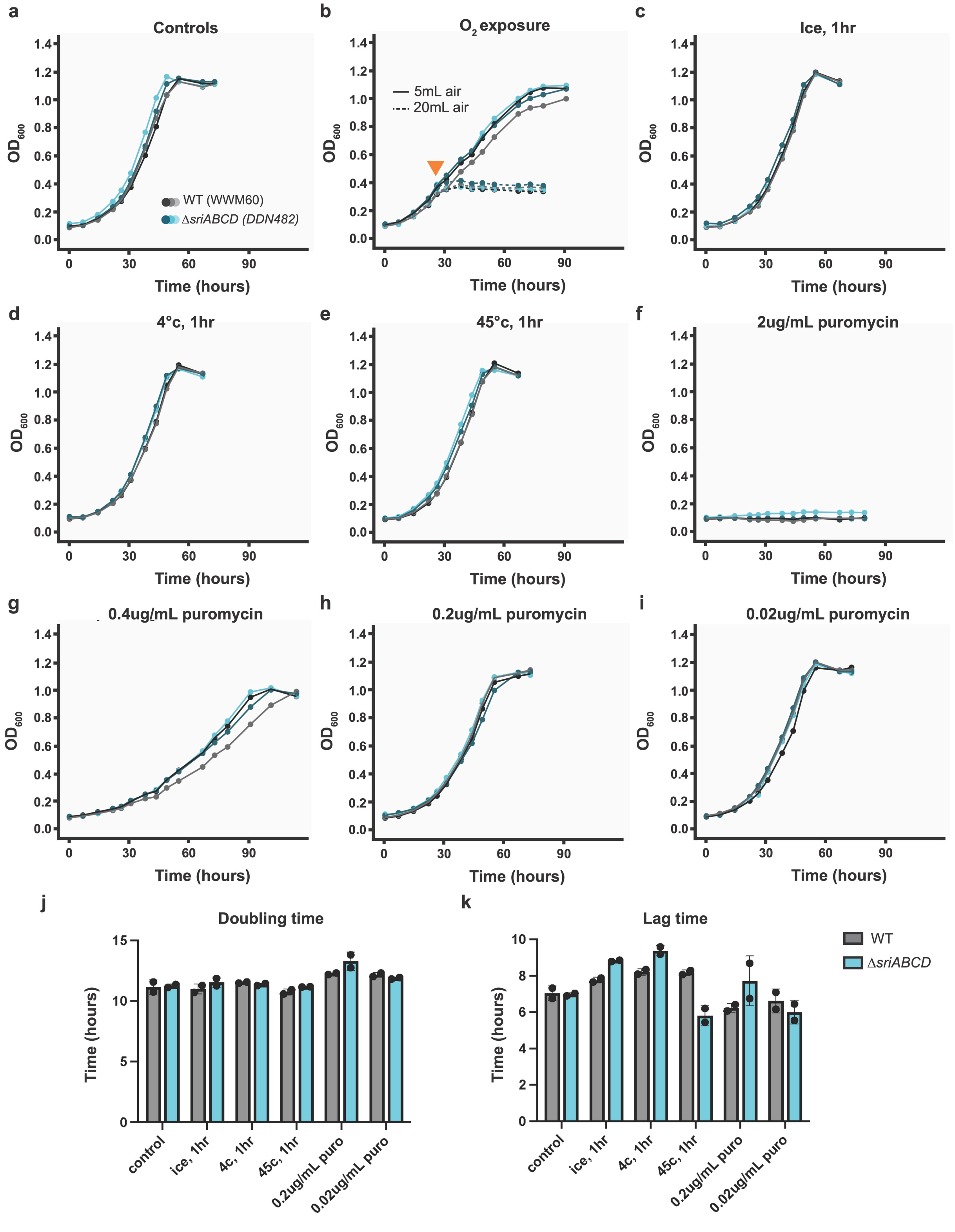
a
Controls
b
O2 exposure
c
Ice, 1hr
5mL air
20mL air
WT (WWM60)
ΔsriABCD (DDN482)
d
4°c, 1hr
e
45°c, 1hr
f
2ug/mL puromycin
g
0.4ug/mL puromycin
h
0.2ug/mL puromycin
i
0.02ug/mL puromycin
OD600
Time (hours)
j
Doubling time
k
Lag time
Time (hours)
WT
ΔsriABCD
control
ice, 1hr
4c, 1hr
45c, 1hr
0.2ug/mL puro
0.02ug/mL puro

**Supplementary Fig. 2:** Challenging Δ*sriABCD* with various stressors yields no growth defect relative to WWM60 (WT). (a-i) WT (WWM60, gray) vs Δ*sriABCD* (blue) growth curves under the following conditions. (a) Controls; standard conditions: 37 °C, anaerobic, 0 μg/mL puromycin. (b) 5 or 20 mL ambient, 0.22 μm filtered air injected into tubes in early exponential phase (indicated by orange carat). (c-e) 1hr incubation immediately after inoculation in ice, 4 °C and 45 °C, respectively; incubated at 37 °C for the remainder of the experiments. (f-i) Growth medium supplemented with 2, 0.4, 0.2, and 0.02 μg/mL puromycin. (j) Doubling and (k) lag times calculated for cultures which grew exponentially (all except b, f, g). Error bars represent standard deviation. All conditions tested in duplicate.

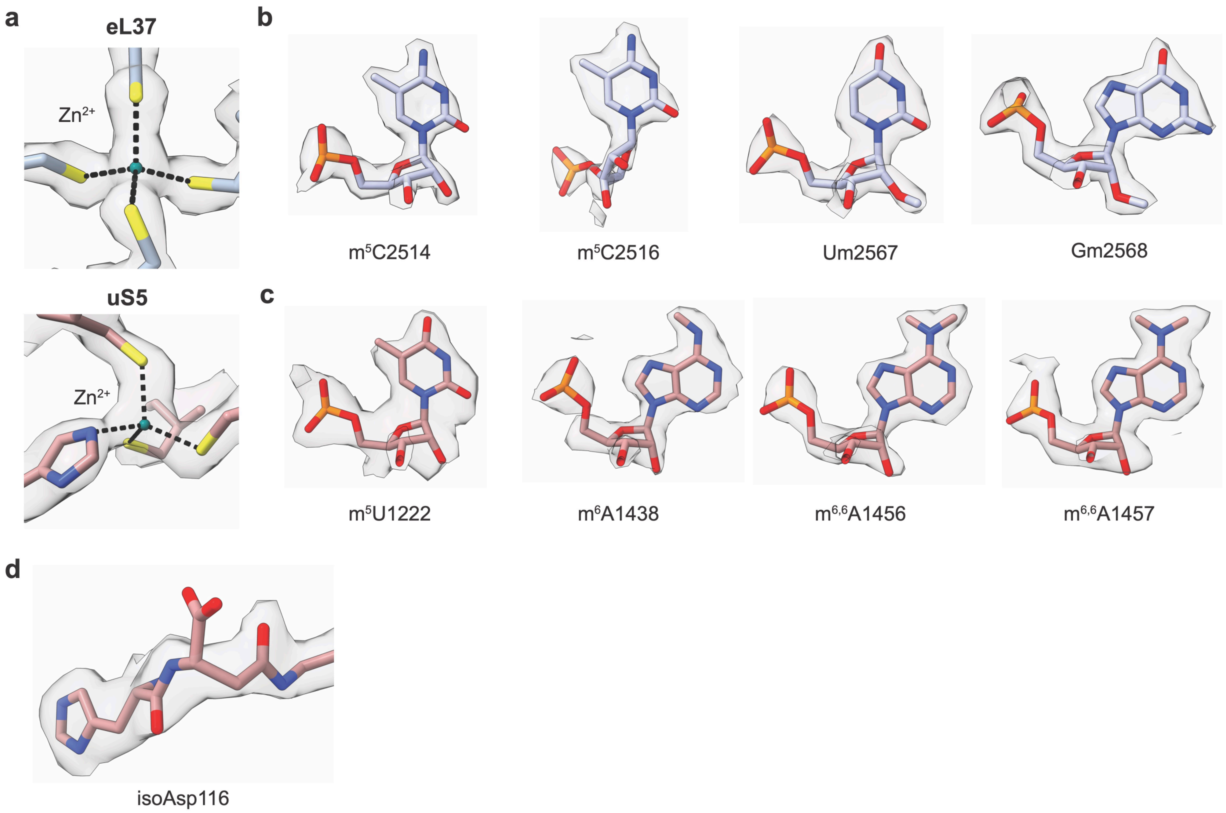


**Supplementary Fig. 3:** Metal binding sites and post-transcriptional and post-translational modifications in the *M. acetivorans* ribosome. (a) Zinc binding sites in LSU rProtein eL37 and SSU rProtein uS5 with cryo-EM density. (b-c) Cryo-EM density for post-transcriptional in the LSU (b) and SSU (c). (d) Cryo-EM density for isoaspartate residue 116 in uS11.

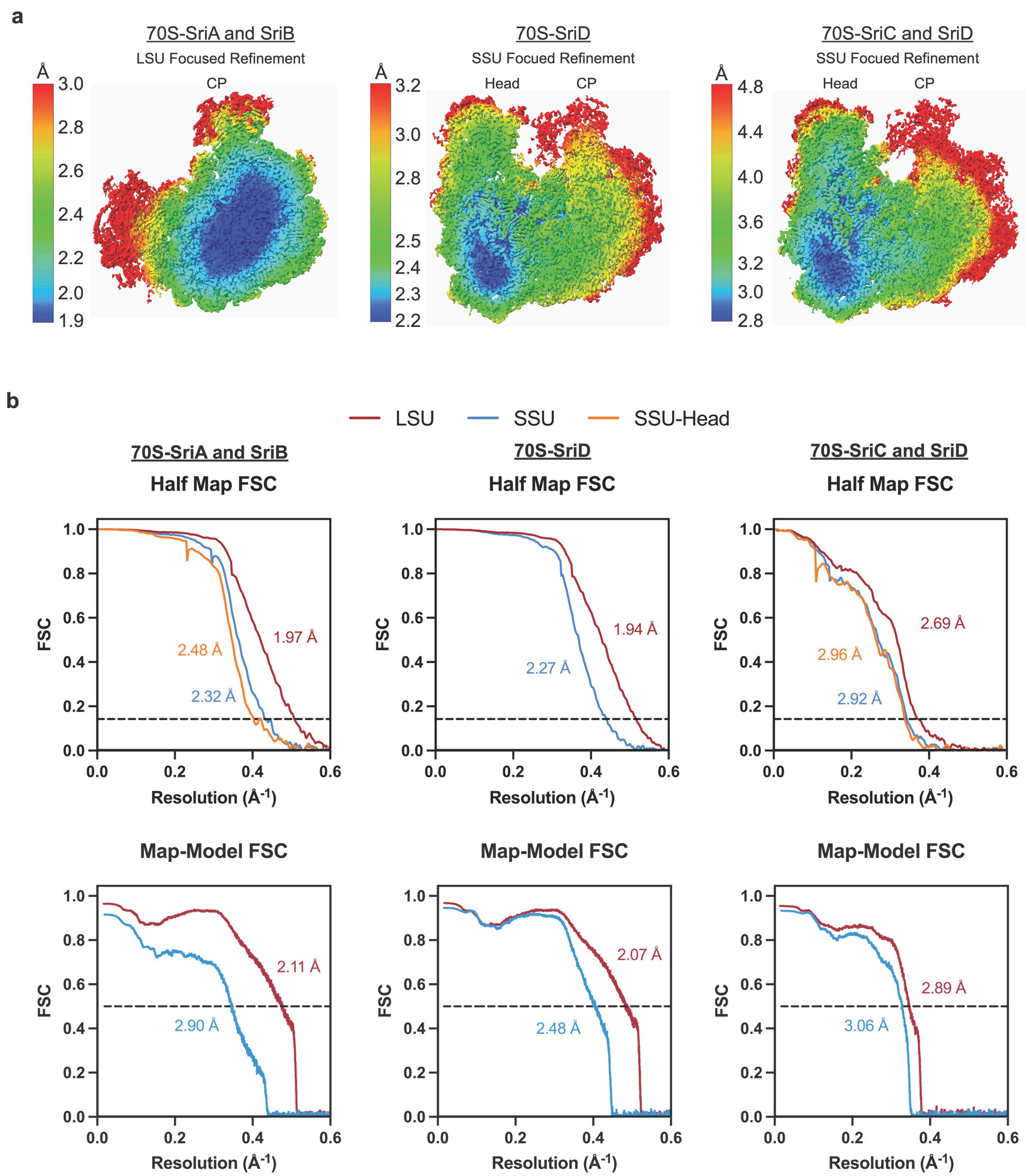


**Supplementary Fig. 4:** Resolution of cryo-EM maps. (a) Local resolution plotted on the LSU (left) or SSU (middle and right) focused refinement maps. The LSU central protuberance (CP) is labelled. (b) Half map (top) and map to model (bottom) FSCs for cryo-EM maps and models in this study.

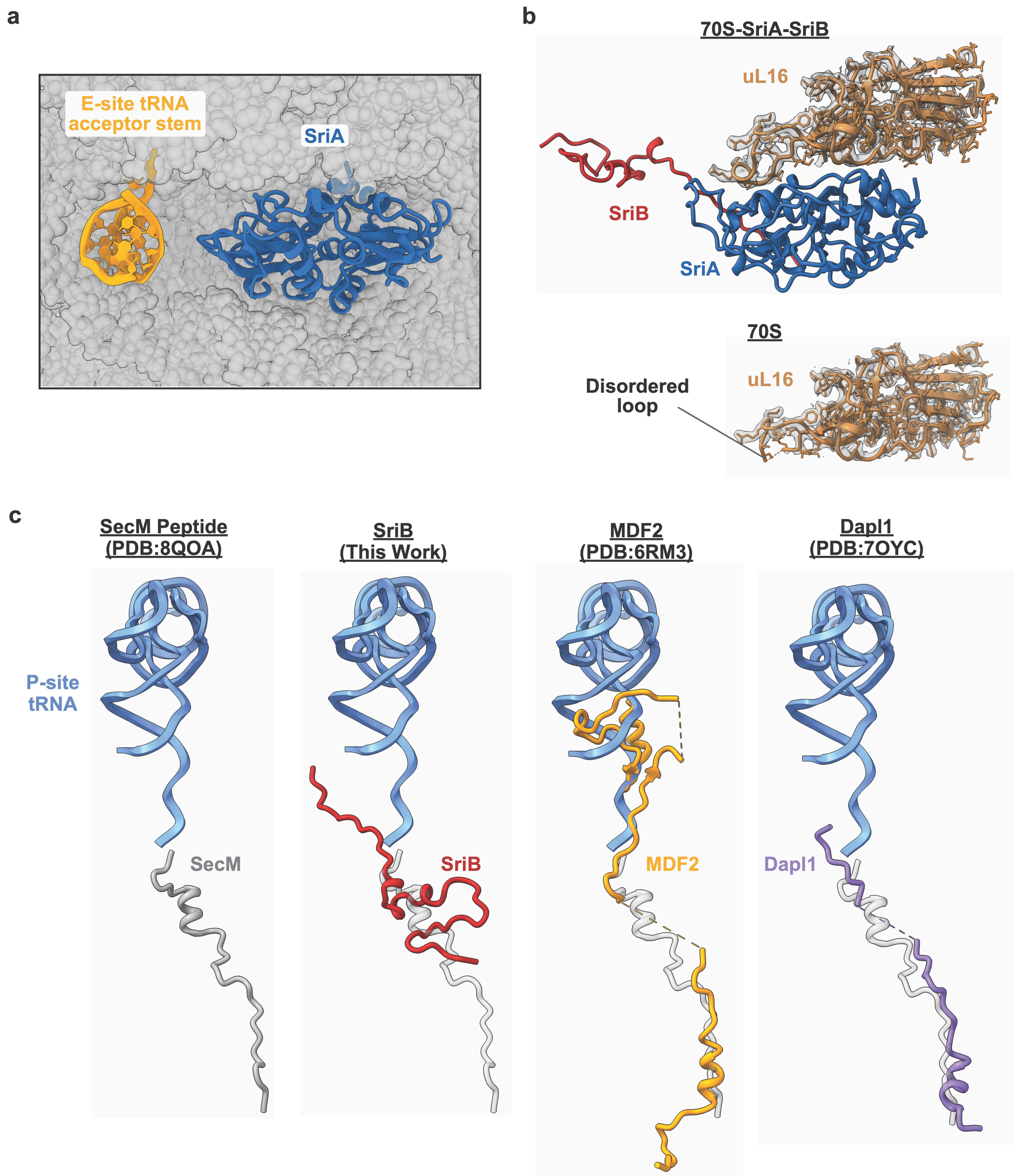


**Supplementary Fig. 5:** SriA and SriB interactions with the LSU. (a) SriA binding is compatible with the presence of E-site tRNA. (b) Model and cryo-EM density of uL16 in the 70S-SriA-SriB complex (top) compared with the 70S ribosome lacking SirA and SriB (bottom). In the absence of SriA and SriB, a loop of rProtein uL16 is disordered. (c)

Comparison of the SriB N-terminus with a stalled nascent polypeptide chain[1] and hibernation factors MDF2[2] and Dapl1[3], which target the exit tunnel. The SecM peptide is overlaid in grey on the hibernation factors to highlight differences in their paths through the PTC and exit tunnel.

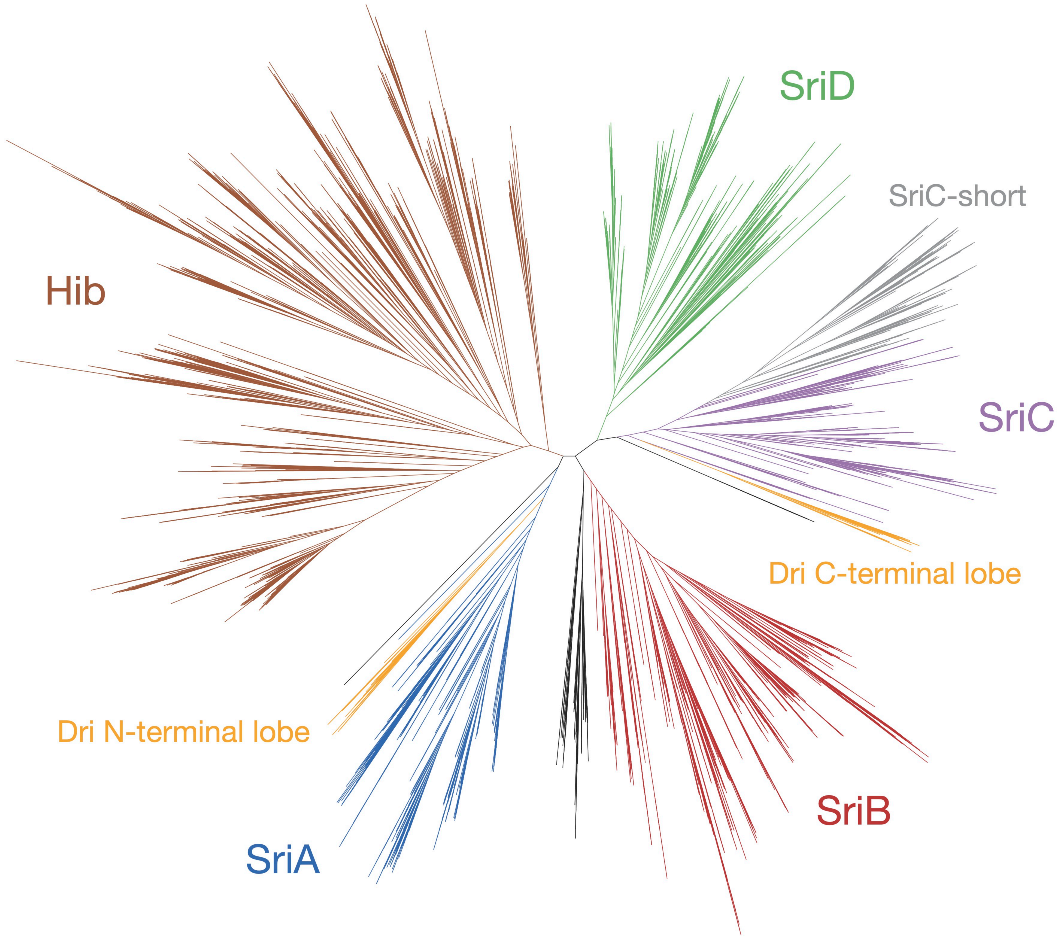


**Supplementary Fig. 6:** Phylogenetic tree of all Sri proteins and related homologs in archaea, including Hib and Dri (split into N- and C-terminal lobes), mined from proteomes of GTDB representative species. Subclades corresponding to SriA, SriB, SriC, SriD, and Hib were defined as regions of the tree that consistently hit the respective homology model best. SriC-short represents a diversification of SriC proteins predominantly in Nitrososphaerales that have a shorter and less positively charged loop that likely function in a different manner than canonical SriC. Clades colored black do not consistently hit one of the homolog models best and may represent related CBS domain-containing proteins.

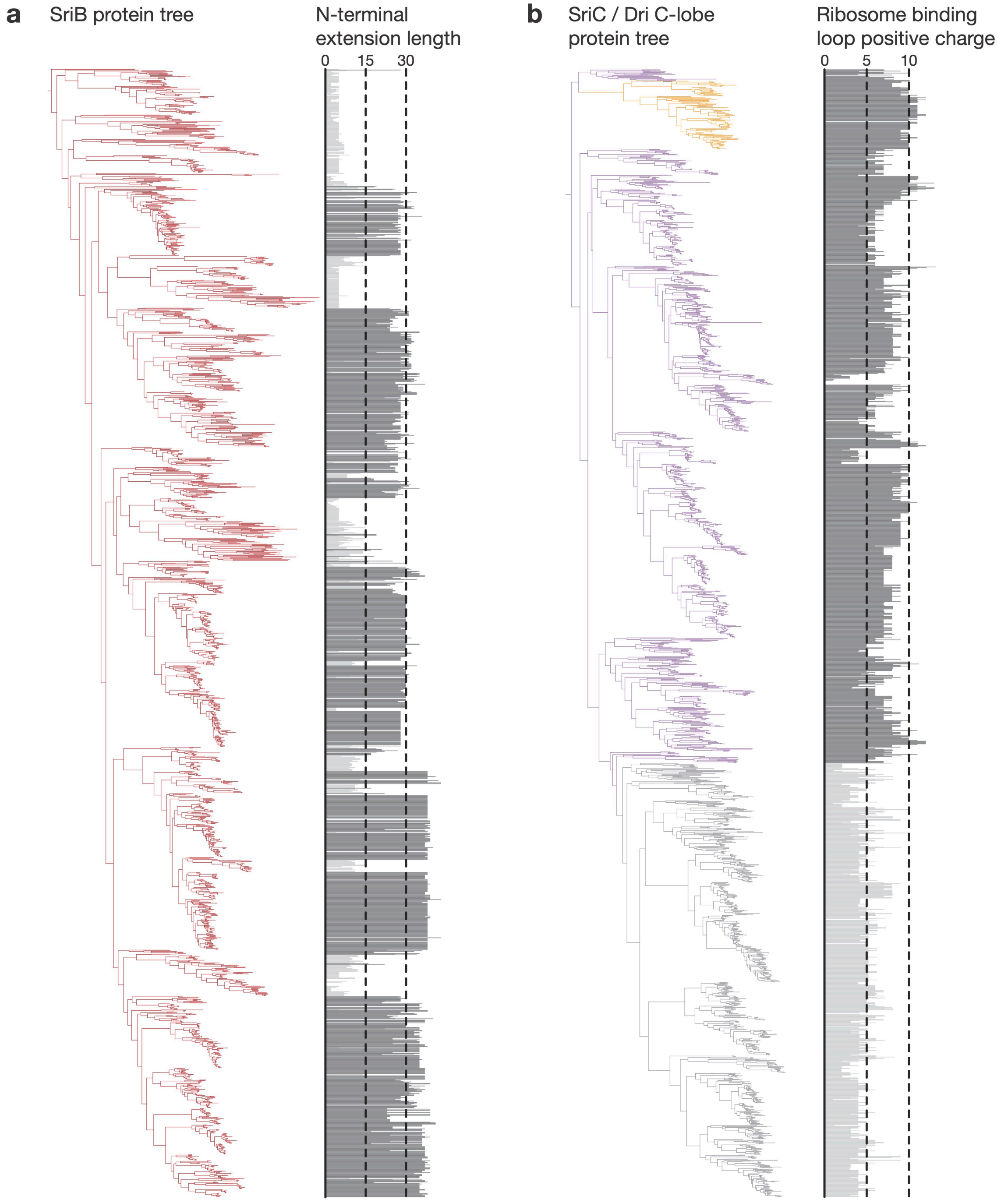


**Supplementary Fig. 7:** SriB and SriC sections of the Sri-Hib protein tree from Supplementary Fig 6. (a) SriB tree with the length of the N-terminal extension plotted for each homolog. Proteins with an extension of less than or equal to 15 amino acids (colored light gray) were designated SriB-short and were considered separately in cooccurence, phylogenetic distribution, and conservation analyses. (b) SriC / Dri C-terminal lobe tree with the number of positively charged residues (adjusted for histidine partial charge) of the ribosome binding loop plotted for each homolog. The subclade highlighted in light gray is named SriC-short, a diversification of SriC proteins predominantly in Nitrososphaerales with on average shorter and less positively charged loops. These sequences were considered separately in co-occurrence, phylogenetic distribution, and conservation analyses.

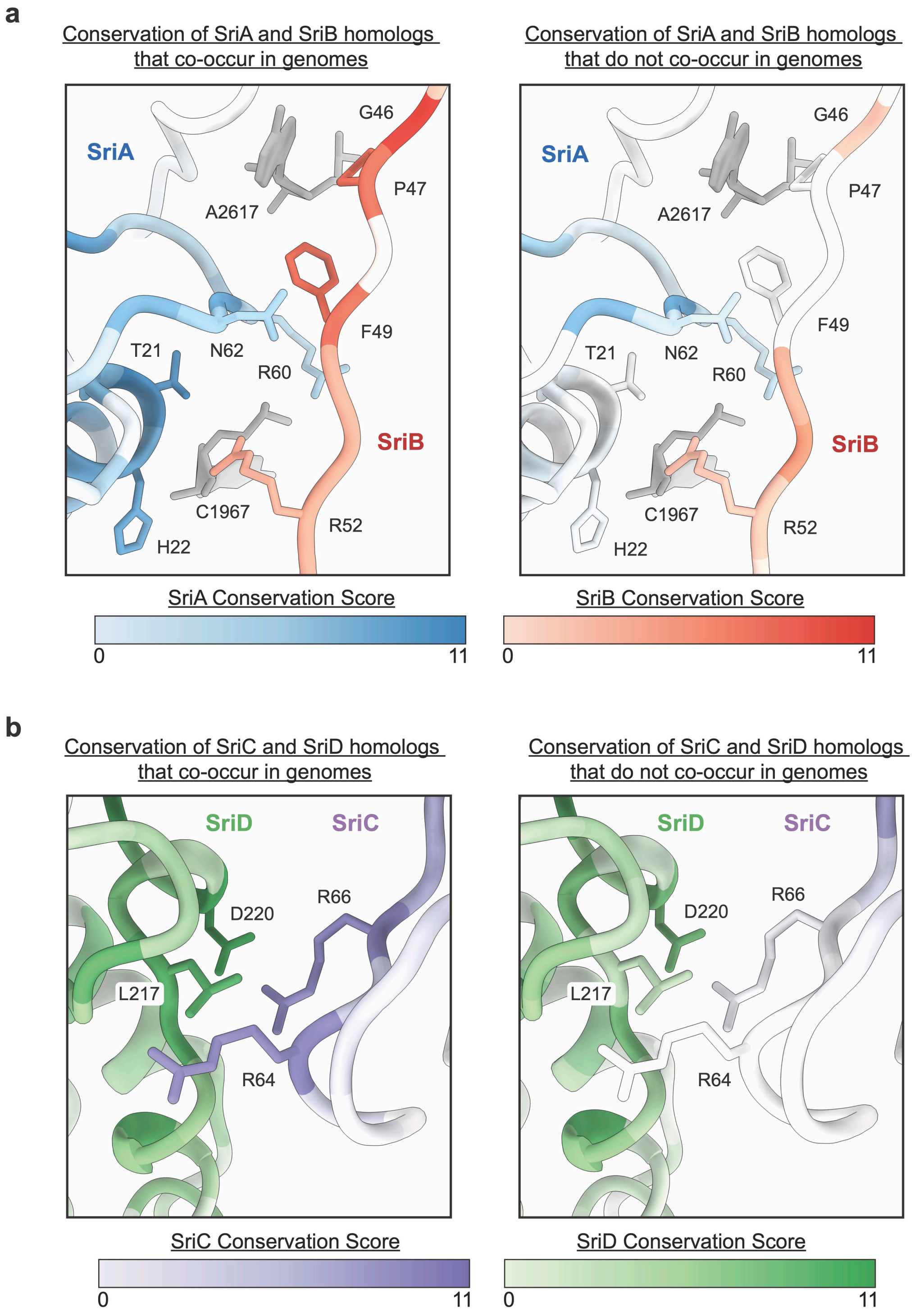


**Supplementary Fig. 8:** Conservation of Sri-Sri interface residues. (a) SriA-SriB interface residues are colored based on conservation scores from homologs in organisms that encode both proteins (left) or in organisms that have standalone *sriA* or *sriB* genes (right). (b) SriC-SriD interface residues are colored based on conservation scores from homologs in organisms that encode both proteins (left) or in organisms that have standalone *sriC* or *sriD* genes (right).

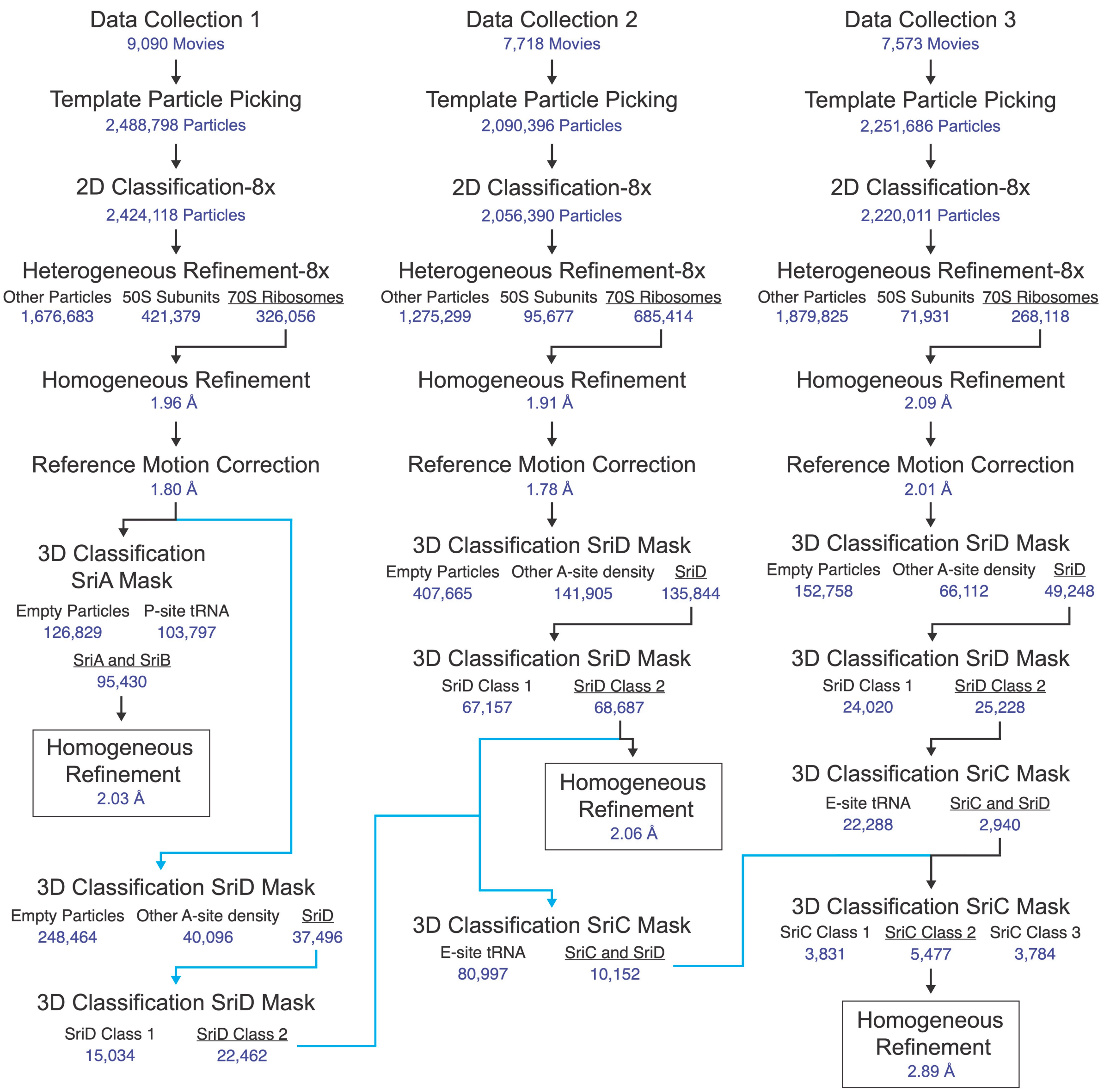


**Supplementary Fig. 9:** Cryo-EM processing workflow. Blue arrows show the processing path to combine SriD containing-particles from the three datasets for SriC classifications and reconstructions.

**Supplementary Table 1. Nucleotide Numbering Conversions**

| ***E. coli*** **23S Numbering** | ***M. acetivorans*** **23S Numbering** | ***E. coli*** **16S Numbering** | ***M. acetivorans*** **16S Numbering** |
|---|---|---|---|
| 1942 | 1967 | 470 | 530 |
| 1944 | 1969 | 1492 | 1430 |
| 2506 | 2521 | 1493 | 1431 |
| 2507 | 2522 | 1494 | 1432 |
| 2583 | 2598 | 1495 | 1433 |
| 2584 | 2599 | 1496 | 1434 |
| 2585 | 2600 | 1534 | 1472 |
| 2586 | 2601 | 1535 | 1473 |
| 2602 | 2617 | | |

**Supplementary Table 2. Primers Used in this Study**

| Name | Sequence | Description |
|---|---|---|
| oMW025 | CTCTTTGATACTTACAGGCCACATGTGCTTACA GCATAATTAGGG | CRISPR guide fragment, MA4648-51 target, R |
| GLC245 | CCTTTTTTTTTCGAAGTTTAACAACATCAGTCAC CTAAAAAG | CRISPR guide fragment with pDN201 overhang, F |
| oMW026 | GGCCTGTAAGTATCAAAGAGGTTTTAGAGCTA GAAATAGCAAGTTAAAATAAG | CRISPR guide fragment, MA4648-51 target, F |
| GLC246 | GCCGGCGCGCCTGCAGGTTTTACATGAGGGC TGAAAAGC | CRISPR guide fragment with pDN201 gibson overhang, R |
| oMW029 | GCCCTCATGTAAAACCTGCAGGGTCAATGGTTG ATGAACCTGAG | MA4648-51 upstream homology gibson overhang, F |
| oMW033 | GAAGTTTATTCCTCGATTGTGCTGACATTCATTG TTACTTTTCCTC | MA4648-51_upstream homology gibson overhang, R |
| oMW034 | AAGTAACAATGAATGTCAGCACAATCGAGGAAT AAACTTCAGAGAAAAG | MA4648-51 downstream homology gibson overhang, F |
| oMW035 | TAAGCGGCCGCGATCGCCGGACGAGATTATTG GGAACGATATGC | MA4648-51 downstream homology gibson overhang, R |
| oMW045 | TTTTTTTAAGATTTCTCGATGTTTTTCATTCTG | *sriABCD* (MA4648-51 region) KO genotype primers, F |
| oMW046 | TGTACTTGAATTTCAATTAAGTTACGAGG | *sriABCD* (MA4648-51 region) KO genotype primers, R |
| oMW047 | GTAATTCCAGACCTTGGAGATCC | *SriABCD* (MA4648-51 region) internal KO genotyping primer, R |
| oMW048 | CCTCAATCTGGAAGAAGAACTGG | *sriABCD* (MA4648-51 region) internal KO genotyping primer, F |
| AB90 | CTCGTAGAAGGGGAGGTTGCG | *pac* (PurR) (for detection of editing plasmid), F |
| AB91 | ATGACCGAGTACAAGCCCACGG | *pac* (PurR) (for detection of editing plasmid), R |

### Supplementary Table 3. Plasmids Used in this Study

| Name | Description | Reference |
|---|---|---|
| pAMG40 | E. coli shuttle vector for fosmid retrofitting to enable replication in *M. acetivorans.* Contains pC2A and λattB | [5] |
| pDN201 | pJK027A derived. Contains PmcrB(tet01)::Spy *cas9* | [6] |
| pMW001 | Gibson assembly of MA4648-50 guide 1 fragment + pDN201 (PmeI) | This study |
| pMW003 | Gibson assembly of pMW001 (AscI) + MA4648-51 homology fragment | This study |
| pMW005 | Co-integrant of pMW003 + pAMG40 | This study |

**Supplementary Table 4. Strains Used in this Study**

| Strain | Genotype | Reference |
|---|---|---|
| WWM60 | *M. acetivorans* ∆*hpt*:: *PmcrB-tetR* | [5] |
| DDN482 | WWM60 ∆*sriABCD* (MA4648-4651) | This study |

**Supplementary Table 5. Protein Sequences Used for Recombinant Expression in this Study**

| Name | Sequence |
|---|---|
| SriA | MNVSEIMSEGPVSIKERDFVTHARQLMRDYLFRSLVVVDEGNRLVGMLNDQDIMRVTS<br>TRSNVTVGGYARPSPTVTPDMDVVKAAKLMVQSKQNRVPVVKSTTDHTVVGVLSDVDI<br>LRNAELPRSASKTIDMVMTKKVKTCSPDERISKVWNYMTETDYTGIPVVSKKGDPIGMI<br>TRRDIIKAGILRMSIEDERAARPNESPKVEKIMSTPAYTLSENDSVKSAIEMIIQHDIGRVT<br>IVNEQGKISGIADRQDLMNAFVNGWSEHHHHHH |
| SriB | MILLNKNTTFVPVEKMNVQPQVNKSGKKAQQKDPHSVSSMGTMRIGPSFKSRIAEHEG<br>KILALATRDVVTLPPTAAIMEAVRIMTERRFRRIPITDAGTGRLEGVVTSVDIIDFLGGGS<br>RNLLVENRFKGNLLAAINEEVRQIMQTDVAYLNDQADFKDAVTTMLERRTGGLPIVNDE<br>MQVIAIFTERNAVELMGGIVTNKTVDEYMTKNVTMVTTDTPIGQAAKVMVQNRFRRLPV<br>VKDGIFAGIVTASDIVHFLGRGDAFSKLTTGNIHEALDQPVGSIVSQELIWTSPGTDMGK<br>AMEIMLEKKIGSLPVLEDGMLRGIITESDFLRGFDLHHHHHH |
| SriC | MNVADIMSSPVYAINIDEPVSRARKLMLRHRISTLLVLNEGKMVGIVTKSDISNRLAQAE<br>PLWRRRPIDQIPIKLLMTESVITIYPEASISQAAALMLENGVHDIPVVKNDIVGIVTRTDIVR<br>YVAEHADEIDTKISTLMTDDIVSVHRHHTINHVIEEMNKNEIERVIVKDDAGKPVGVISKR<br>NLALNLLTDNEGKLSTKSIKMARKSSPGGQKTYRYVKEVPLTAEDIMITPIISIDVNEKISI<br>AAKKLIEEEITALPVSDGEEIVGILSRTDIMKSVLHHHHHH |
| SriD | MQRIETKTLDTHEVKCMQVKDIMVQPHKIDKSDTISHALDLMEKKDTKRLLVVHDNQVL<br>GVLTMRGLTEQLGTRRKQSKPASSLHVATAVSDNFVKVLPDTDVKDALTLMKKKGGVII<br>VTDNGNAMGWVTPQELMKVNHFTGFAGEVMEKNPIIVSPSDRVSHARRLILDKNVGRL<br>PVIENGKLVGIIAEDDIAFAMRSFRDLVADNQQDSRIKNLLVGDIMTRSVVNVYTNTPLS<br>DTVDTMLEYDVGGVPVLNLEEELVGFLARRNIINTIEEHHHHHH |

### Supplementary Table 6. NLuc *In Vitro* Translation Reporter mRNA Sequence

The SD sequence is **bolded**; The start codon is underlined

| Name | Sequence |
|---|---|
| NLuc mRNA | GGUCCCAAAGUGAUUUAAUAAAUU**AAAGGAGGA**AAUUAAAAUGGUCUUCACACU<br>CGAAGAUUUCGUUGGGGACUGGCGACAGACAGCCGGCUACAACCUGGACCAAGU<br>CCUUGAACAGGGAGGUGUGUCCAGUUUGUUUCAGAAUCUCGGGGUGUCCGUAA<br>CUCCGAUCCAAAGGAUUGUCCUGAGCGGUGAAAAUGGGCUGAAGAUCGACAUCC<br>AUGUCAUCAUCCCGUAUGAAGGUCUGAGCGGCGACCAAAUGGGCCAGAUCGAAA<br>AAAUUUUUAAGGUGGUGUACCCUGUGGAUGAUCAUCACUUUAAGGUGAUCCUGC<br>ACUAUGGCACACUGGUAAUCGACGGGGUUACGCCGAACAUGAUCGACUAUUUCG<br>GACGGCCGUAUGAAGGCAUCGCCGUGUUCGACGGCAAAAAGAUCACUGUAACAG<br>GGACCCUGUGGAACGGCAACAAAAUUAUCGACGAGCGCCUGAUCAACCCCGACG<br>GCUCCCUGCUGUUCCGAGUAACCAUCAACGGAGUGACCGGCUGGCGGCUGUGC<br>GAACGCAUUCUGGCGUAAGGAUCCGGAGAGCUCCCAACGCGUUGGAUGCAUAG<br>CUUGAGUAUUC |

### Supplementary Table 7. Cryo-EM Data Collection and Processing

| **Reconstruction** | **70S-SriA-SriB** | **70S-SriD** | **70S-SriC-SriD** |
|---|---|---|---|
| Magnification | 105,000 | 105,000 | 105,000 |
| Voltage (kV) | 300 | 300 | 300 |
| Electron Exposure ($e^-$/Å$^2$) | 40 | 40 | 40 |
| Defocus Range (μm) | -0.5/-1.5 | -0.5/-1.5 | -0.5/-1.5 |
| Pixel Size (Å) | 0.8264 | 0.8264 | 0.8264 |
| Symmetry Imposed | C1 | C1 | C1 |
| Initial Particle Images | 2,488,798 | 2,090,390 | 6,830,880 |
| Final Particle Images | 95,430 | 68,687 | 5,477 |
| Map Resolution (Å) | 2.03 | 2.06 | 2.89 |
| FSC Threshold | 0.143 | 0.143 | 0.143 |

### Supplementary Table 8. Model Refinement Statistics

| Model | **70S-SriA-SriB** | **70S-SriD** | **70S-SriC-SriD** |
|---|---|---|---|
| Model component | | | |
| Model resolution (Å) | 2.11/2.90 (50S/30S) | 2.07/2.48 (50S/30S) | 2.89/3.06 (50S/30S) |
| FSC threshold | 0.5 | 0.5 | 0.5 |
| Map sharpening *B* factor (Å$^2$) | 35.5/48.6 (50S/30S) | 30.7/48.3 (50S/30S) | 25.1/27.9 (50S/30S) |
| Model composition | | | |
| Non-hydrogen atoms | 157707 | 159420 | 150576 |
| $Mg^{2+}$ ions | 305 | 314 | 314 |
| Waters | 8740 | 10476 | 0 |
| Mean *B* Factors (Å$^2$) | | | |
| RNA | 51.58 | 49.86 | 44.19 |
| Protein | 62.40 | 60.01 | 51.40 |
| Waters | 50.41 | 48.73 | - |
| Other | 58.21 | 56.95 | 33.33 |
| R.m.s. deviations from ideal values | | | |
| Bond (Å) | 0.005 | 0.005 | 0.006 |
| Angle (°) | 0.655 | 0.669 | 0.843 |
| Molprobity score | 1.86 | 1.89 | 2.26 |
| Clash Score | 7.91 | 7.50 | 8.60 |
| Rotamer outliers (%) | 2.59 | 2.82 | 5.04 |
| Ramachandran plot | | | |
| Favored (%) | 97.39 | 97.28 | 96.14 |
| Allowed (%) | 2.50 | 2.68 | 3.77 |
| Outliers (%) | 0.11 | 0.04 | 0.09 |
| RNA validation | | | |
| Angles outliers (%) | 0.01 | 0.01 | 0.01 |
| Sugar pucker outliers (%) | 0.9 | 0.7 | 0.7 |
| Average suiteness | 0.634 | 0.633 | 0.596 |
